# Intracellular sphingolipid sorting drives membrane phase separation in the yeast vacuole

**DOI:** 10.1101/2023.07.13.548923

**Authors:** Hyesoo Kim, Itay Budin

## Abstract

The yeast vacuole membrane can phase separate into ordered and disordered domains, a phenomenon that is required for micro-lipophagy under nutrient limitation. Here we report that sorting of sphingolipids (SLs) into the vacuole membrane controls this process. We first developed a vacuole isolation method to identify lipidome changes during the onset of phase separation in early stationary stage cells. We found that phase separated vacuoles are characterized by increases in lipid raft-forming components not found in the whole cell, including a dramatic change in SL composition. Sorting of both SLs and ergosterol into the vacuole membrane is dependent on Npc2, the yeast homologue of the Niemann-Pick Type C2 lipid transporter. Genetic dissection of SL biosynthesis revealed that the composition of vacuole SLs modulates membrane phase separation and micro-lipophagy under glucose restriction. These results show that lipid trafficking can drive membrane phase separation *in vivo* and identify SLs as key mediators of this process in yeast.

## Introduction

The ability of lipid-lipid interactions to drive heterogeneity within continuous bilayers has been proposed as a mechanism for organization in cell membranes. The lipid raft hypothesis postulates that biological membranes can phase separate into coexisting liquid-ordered (Lo) and liquid-disordered (Ld) domains based on lipid composition and temperatures (Owen et al., 2012). Sterols, such as cholesterol in metazoans or ergosterol in fungi, and lipids with saturated acyl chains, including sphingolipids (SLs), are required to form Lo domains and can thus be referred to as raft components. In contrast, Ld domains are enriched in unsaturated glycerophospholipids (GPLs) (Simons and Ikonen, 1997). The formation of Lo domains has been suggested as a sorting mechanism that sequesters raft proteins, such as those with long transmembrane domains or GPI anchors (Sangiorgio et al., 2004). Lipid rafts were first proposed based off of co-segregation of lipids and proteins in detergent resistant fractions of apical epithelial membranes (Schuck et al., 2003) and later characterized extensively in synthetic giant unilamellar vesicles (GUVs) composed of ternary mixtures of lipids (Veatch and Keller, 2003). However, large Lo membrane domains are not readily observable in mammalian cells (Munro, 2003) and current models for lipid rafts focus on small (<100 nm), dynamic assemblies only accessible by superresolution imaging approaches (Hancock, 2006; Levental et al., 2020; Shelby et al., 2023).

While initial work on membrane domains has focused on the mammalian plasma membrane (PM), the vacuole of budding yeast (*Saccharomyces cerevisiae*) has become a surprising and robust model for micron-scale membrane phase separation. Vacuoles are digestive organelles similar to mammalian lysosomes (Thumm, 2000) and their membrane organization has been implicated in microautophagy-related degradation of energy stores during nutritional starvation (Rahman et al., 2021). Freeze fracture electron microscopy studies first suggested that vacuole membranes can display lateral heterogeneity that is dependent on the growth stage (Moor and Mühlethaler, 1963; Moeller and Thomson, 1979; Moeller et al., 1981). More recently, this phenomenon has been further explored using fluorescence microscopy. In exponentially growing cells, vacuole membrane-associated proteins expressed as fluorescent protein fusions are evenly distributed across the surface, but in stationary phase segregate into discrete patterns of domains and surrounding areas (Toulmay and Prinz, 2013). Vacuole domains, labeled with Lo proteins such as the BAR-domain containing protein Ivy1, are often polygonal in shape and are stained with the sterol-binding dye fillipin. In contrast, the surrounding areas are labeled with Ld proteins, such as the vacuolar ATPase Vph1 or alkaline phosphatase Pho8 (Leveille et al., 2022), and are stained with lipophilic dyes that localize to Ld domains in GUVs, like FAST DiI. Vacuole domains dissolve above a characteristic lipid melting temperature and this process is reversible (Rayermann et al., 2017), similar to domain formation in phase separated GUVs. In cells, the formation of vacuole Lo domains is dependent on autophagy related machinery, and they serve as docking sites for lipid droplet (LD) internalization (Seo et al., 2017). Once in the vacuole, LDs are digested to release fatty acids that drive energy metabolism via β-oxidation in the mitochondria during conditions of glucose deprivation, termed micro-lipophagy (Rambold et al., 2015). Cells lacking vacuole domains are defective in micro-lipophagy and show poor survival under nutritional stress, while those lacking autophagy machinery, like Atg14, fail to form vacuole domains (Seo et al., 2017), suggesting these processes in tightly linked.

Although biological functions for vacuole domains have been identified, the specific changes in membrane lipid or protein composition that drive their formation have not yet been defined. Sterols, which are predominantly in the form of ergosterol in fungi, are hallmarks of Lo domains and are required for their formation *in vitro* (Veatch and Keller, 2003). Yeast vacuole domains are lost upon ergosterol extraction by methyl-β-cyclodextrin and fail to form when cells are grown with the ergosterol biosynthesis inhibitor fenpropimorph (Toulmay and Prinz, 2013). Reduction in whole cell ergosterol levels through repression of sterol biosynthesis similarly also modulates vacuole domain abundance and size (Seo et al., 2021). Sterols could be trafficked to the vacuole membrane either from organelles via vesicular pathways, such as the vacuole fusion of multivesicular bodies (MVB) (Schulz and Prinz, 2007), or non-vesicular machinery, such as through the Niemann-Pick type C (NPC) homologs Ncr1 and Npc2 (Tsuji et al., 2017; Leveille et al., 2022). Loss of NPC proteins reduces the frequency of membrane domains (Tsuji et al., 2017), suggesting that lipid trafficking to the vacuole membrane could be involved in phase separation. Unlike its mammalian counterpart, Npc2 has a large binding cavity (Winkler et al., 2019) that can accommodate a range of non-sterol substrates and *in vitro* binds polar lipids, including anionic and inositol-containing GPLs (Moesgaard et al., 2020).

To form Lo domains, sterols must preferentially interact with saturated lipids, which in eukaryotic cells are predominantly SLs. In contrast to metazoans, complex sphingolipids (CSLs) in yeast are anionic and contain one of three head groups composed of inositol phosphate and mannose. The resulting CSLs – inositol phosphorylceramide (IPC), mannosyl inositol phosphorylceramide (MIPC), and mannosyl diinositol phosphorylceramide (M(IP)_2_C) – are among the most abundant lipids in yeast cells, together comprising 10-15 mol % of the lipidome (Ejsing et al., 2009). Overall CSL concentration stays constant during yeast growth stages, but the stoichiometry between the three species changes (Klose et al., 2012). CSLs are formed from a ceramide (either phytoceramides or dihydroceramide) backbone that feature very long chains, with a combined length of up to 46 carbons, that can be additionally hydroxylated (Ejsing et al., 2009). During SL metabolism, phytoceramides or dihydroceramides are first synthesized in the ER, transferred to Golgi for modification, and then sorted for transport to other compartments (Levine et al., 2000; Funato and Riezman, 2001; Olson et al., 2016).

It has been proposed that domain formation in stationary stage or starved cells occurs via a redistribution of raft-forming lipids within the cell, leading to an increase in their abundance under these conditions (Seo et al., 2021). Directly testing this model has been hindered by the challenges in isolating stationary stage vacuoles, including poor cell wall lysis and growth stage differences in vacuole density. Recently, a method for immunoprecipitation of vesicles derived from yeast organelles has been developed, termed MemPrep (Reinhard et al., 2022), and has been applied to the analysis of stationary vs. exponential stage vacuoles sampled at 48 and 8 hours of growth, respectively (Reinhard et al., 2023). Stationary stage vacuole membranes isolated by MemPrep using the bait vacuole protein Mam3, an Ld marker, showed no increase in ergosterol content, and overall low levels of both ergosterol and sphingolipids. Instead, an increase in the abundance of high melting temperature phosphatidylcholine (PC) species was observed, which also increased in the corresponding whole cell lipidome.

In this study, we test the hypothesis that redistribution of raft-forming lipids into the vacuole drives its membrane to phase separate under glucose restriction. We find that the lipidomes of early stationary stage vacuoles isolated by density centrifugation show a clear enrichment in Lo-domain promoting lipids, especially CSLs that increase several-fold in abundance compared to late exponential stage vacuoles. These increases are not found in the corresponding whole cells and are dependent on yeast homologues of NPC proteins. We then perform a systematic genetic analysis of the yeast SL pathway to dissect how the SL composition controls microdomain abundance, morphology, and microautophagy of LDs during nutritional stress.

## Results

### Lipidomic analysis of isolated yeast vacuoles under domain-forming conditions

We first defined growth conditions in which liquid cultures showed robust phase separated vacuoles at stationary stage, as imaged by the well-established Ld marker Pho8-GFP (Fig. 1A). Cells in early stationary stage, after 24 hours of growth in minimal medium, showed robust vacuole phase separation but still had cell walls that were readily digestible, unlike those incubated for longer time periods or grown under more complete media. Cells in the late exponential stage, after 17 hours of growth, contained vacuoles of similar size to stationary stage cells, which is ideal for utilizing the same density centrifugation protocol, but has no detectable domains. We then optimized a set of high and low speed density centrifugation steps (Fig. 1B) to generate homogenous vacuole populations that were free of other organelles and LDs. A key addition was a final low speed centrifugation step after the two high speed ones to remove bound LDs. Initial assaying of purity was done by microscopy and then validated by western blotting against vacuole (Pho8) and non-vacuole (mitochondria, Cox4; ER Dpm1; LD, Erg6) proteins (Fig. 1C).

**Figure 1:**
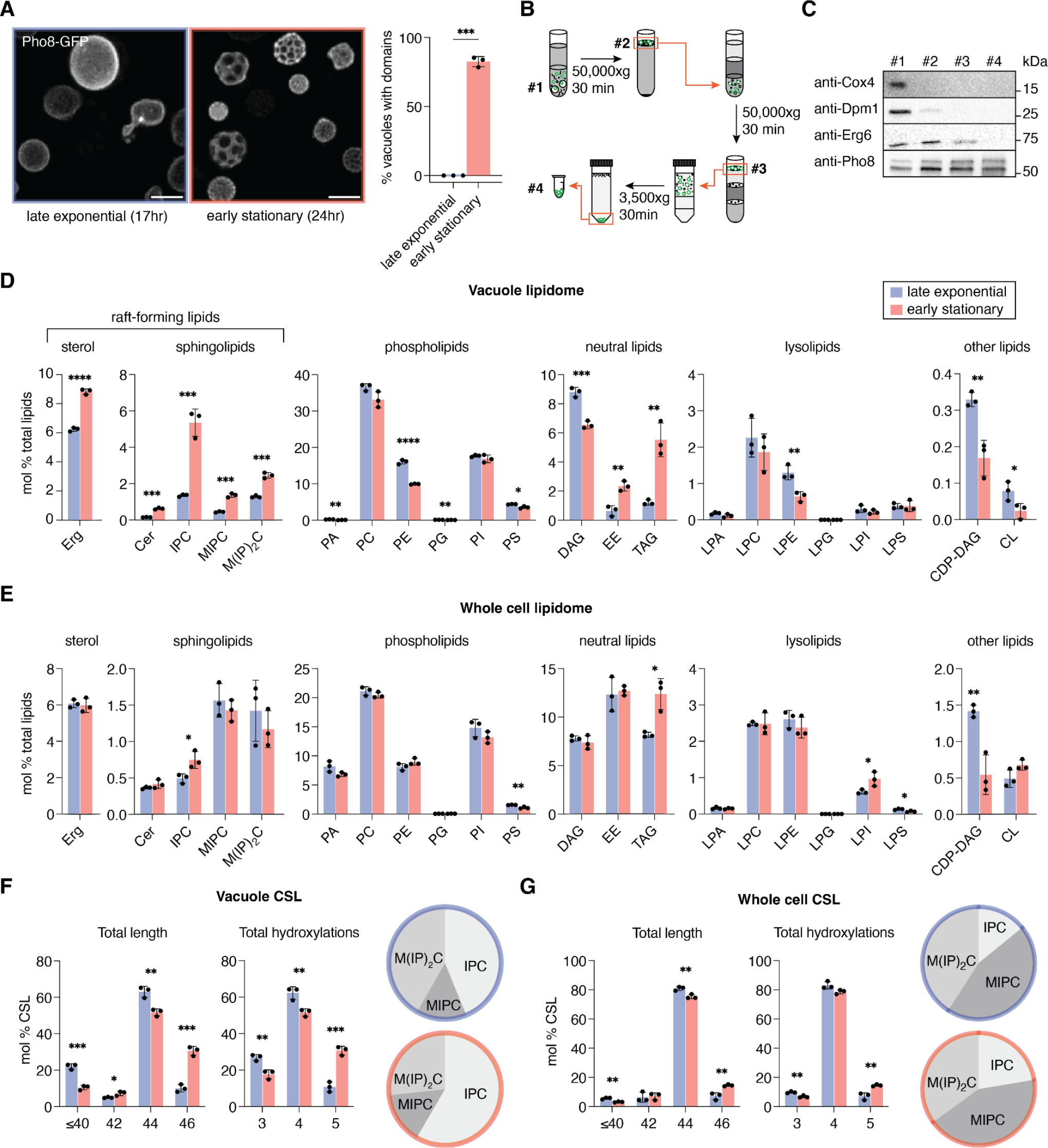
Sorting of raft-forming lipids into phase separated yeast vacuoles. (**A**) Vacuole membranes phase separate into discrete domains during a short growth period (7 hours) as cells enter the stationary stage. 3D projections shown are from confocal Z-stacks of a field of live cells expressing the vacuole Ld domain marker Pho8-GFP grown under vacuole isolation conditions. Scale bar, 3 μm. The bar graph shows the fraction of cells grown under early stationary stage isolation conditions show phase separation, while none do under the late exponential stage conditions. Error bars represent SD for n=3 individual cultures; N > 100 cells per replicate. (**B**) Centrifugation-based isolation of stationary stage vacuoles. Vacuoles were separated from cell lysates via two rounds of density gradient ultracentrifugation, followed by a low-speed centrifugation step to remove associated LDs. (**C**) Western blot analysis of the indicated fractions in the diagram shows how the purification method successively depletes non-vacuole components (Cox4, mitochondria; Dpm1, ER; Erg6, LDs) from stationary stage vacuoles. (**D**) The abundance, expressed as a mol % of total lipids, for each lipid class in vacuoles isolated from W303a cells in late-exponential stage (17 hours of growth, blue) and early stationary stage (24 hours of growth, red) in minimal medium. Error bars indicated SD for n=3 replicate preparations from individually grown cultures. Significance was assessed by unpaired two-tailed t-test; *, p < 0.05; **, p < 0.01; ***, p <0.001. Abbreviations: CDP-DAG, cytidine diphosphate-diacylglycerol; Cer, ceramide; DAG, diacylglycerol; Erg, ergosterol; LPA, lysophosphatidic acid; LPC, lysophosphatidylcholine; LPE, lysophosphatidylethanolamine; LPG; lysophosphatidylglycerol; LPI, lysophosphatidylinositol; LPS, lysophosphatidylserine; PG, phosphatidylglycerol; PI, phosphatidylinositol (**E**) The corresponding lipidomes from intact whole cells grown identically as for vacuole purification, which do not show increases in raft-forming lipids. (**F**) Changes in the CSL pool from late exponential to early stationary stage vacuoles, including, lengthening of total chain length, and increase in the number of hydroxylations, and changes in relative headgroup composition displayed as a pie-chart. ****, p <0.0001. (**G**) The corresponding data for the CSL pool in whole cell lipidomes, which shows smaller changes from late exponential to early stationary stage cells.

We next carried out comprehensive lipidomic analysis of polar and non-polar lipids from purified exponential and stationary stage vacuoles (Fig. 1D), as well as corresponding whole cell samples (Fig. 1E). Distinct features of the vacuole lipidome, such as the low abundance of phosphatidic acid (PA), were consistent between exponential stage vacuoles in our data set, those generated by MemPrep (Reinhard et al., 2023, 2022), and earlier analyses that were also performed by density centrifugation (Zinser and Daum, 1995). Exponential and stationary stage vacuoles isolated by our protocol contained low amounts (<0.1 %) of cardiolipin (CL), a marker of mitochondrial contamination, less than those in MemPrep-isolated vacuoles (Reinhard et al., 2022) or in previously published vacuole lipidomes (Tuller et al., 1999). Because vacuoles form contacts with LDs for degradation of storage lipids, including ergosterol esters (EE) and triacylglycerides (TAG) (Leber et al., 1994), LD contamination is especially challenging for vacuole isolates. Purified vacuoles showed lower levels of neutral lipids – 0.6 mol% EE and 1.2 mol% TAG in exponential vacuoles and 2.3% EE and 5.5% TAG in stationary vacuoles – than previously reported.

### Phase-separated vacuoles are characterized by an enrichment of raft-forming lipids

The lipidome of centrifugation-purified stationary stage vacuoles from W303a (WT) showed a substantial increase of putative raft-forming lipids, including both ergosterol and SLs (Fig. 1D). Compared to late exponential stage vacuoles, the amount of ergosterol increased by 40% and the amount of CSLs showed a nearly 3-fold increase (3.5±0.1 to 10.7±1.0 % of all polar lipids). While all three of the major CSLs showed significant increases, the largest was in IPC (4.0-fold). Thus, the distribution of the CSL pool shifted from one with equal amounts of IPC and M(IP)_2_C in exponential stage, as previously reported (Zinser and Daum, 1995), to one dominated by IPC in early stationary stage vacuoles (Fig. 1F). In the corresponding whole cell lipidomes, the amount of ergosterol, MIPC, and M(IP)_2_C did not increase from late exponential to early stationary stage (Fig. 1E). IPC did show a modest increase, but to a much smaller extent (1.6-fold) than in the vacuole, and the overall composition of SL headgroups were similar between the two growth stages (Fig. 1G). In the vacuole, the average carbon chain length and number of hydroxylation of CSL chains also increased substantially in stationary phase vacuole (Fig. 1F). These changes were also present in the whole cell lipidome, as previously observed (Klose et al., 2012), but to a smaller extent (Fig. 1G). These results indicate that raft-forming lipids are sorted into the vacuole membrane during stationary stage growth, potentially driving phase separation.

We also observed changes in GPLs of stationary stage vacuoles (Fig. 1D), though these were more modest than those for sterols and SLs. Phosphatidylethanolamine (PE) decreased in stationary stage vacuoles, which was also observed in MemPrep-isolated vacuoles (Reinhard et al., 2023), while it slightly increased in the whole cell. However, we did not observe a corresponding increase in phosphatidylserine (PS) or the substantial increase in PC that was previously reported in late stationary phase vacuoles. As discussed further below, these discrepancies could result from the growth stage analyzed, since they are also reflected in the whole cell lipidome data (Fig. 1E). We could also observe changes to the acyl chains of major vacuole GPLs as yeast enter stationary stage (Fig. S1), which were also observed in whole cell lipidomes (Fig. S2). Overall, the PL composition of both late exponential stage vacuoles in our growth conditions were similar to those reported in earlier studies (Zinser and Daum, 1995).

### Sorting of SLs into stationary stage vacuoles is dependent on the lipid transporter Npc2

We next asked by what mechanism raft-promoting lipids sort into the vacuole membrane during the early stationary stage. We focused on the yeast homologues of Niemann-Pick Type C proteins, vacuole lipid transporters whose loss has previously been observed to reduce membrane phase separation (Tsuji et al., 2017). Consistent with previous experiments, we observed that *npc2*Δ cells largely lost the ability to form vacuole domains, while *ncr1*Δ cells showed only a small loss in domain formation (Fig. 2A). We then purified early stationary stage vacuoles from both strains and carried out lipidomic analysis. Compared to WT, early stationary stage vacuoles from *npc2*Δ cells contained lower amounts of all SLs and ergosterol (Fig. 2B). In contrast, *ncr1*Δ vacuoles showed a redistribution of CSL species – an increase in MIPC at the expense of M(IP)_2_C and IPC – but no overall change in their total levels. In the whole cell lipidomes, neither *ncr1*Δ nor *npc2*Δ showed altered CSL levels (Fig. 2C), though we did detect decrease in specific ceramides that were previously observed for these mutants (Vilaça et al., 2018) (Fig. S3 A). There were also subtle changes in the GPL content (Fig. S3 B) of the vacuole, an increase in phosphatidylinositol (PI) and loss of PC, but not in the whole cell (Fig. S3 C). These data suggest that Npc2, but not necessarily its canonical receptor Ncr1, plays a role in vacuole sorting of raft-forming lipids at the onset of membrane phase separation (Fig. 2D).

**Figure 2:**
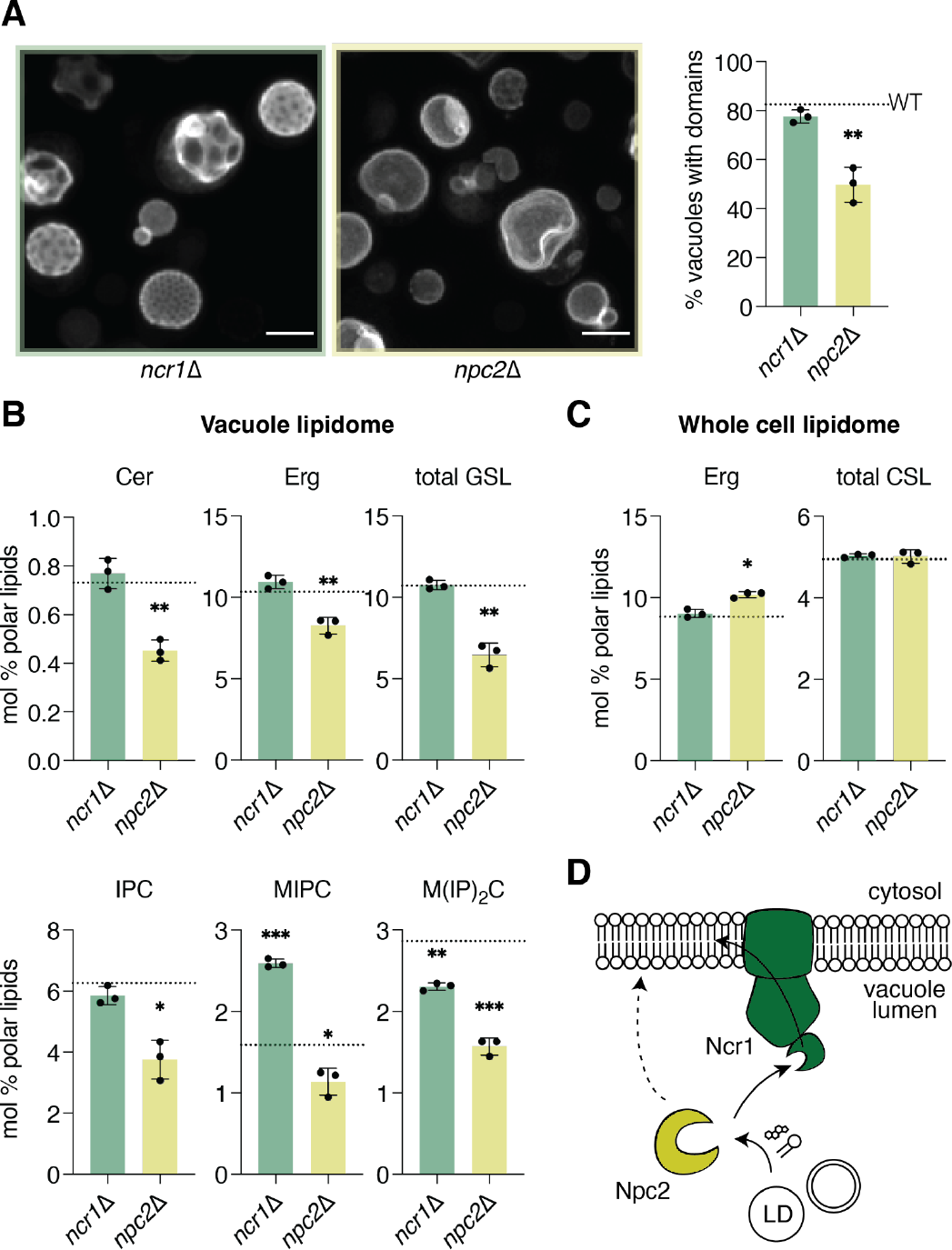
The levels of raft-forming lipids in stationary stage vacuole is dependent on Npc2. (**A**) 3D projections showing a field of early stationary stage cells expressing the vacuole Ld domain marker Pho8-GFP grown under vacuole isolation conditions. Quantification shows that *npc2*Δ cells lose domain formation capability compared to WT cells (dashed line), as previously observed. Significance was assessed by unpaired two-tailed t-test against WT; *, p < 0.05. (**B**) Abundance of raft components in early stationary stage vacuoles isolated from *npc2*Δ and *ncr1*Δ cells. Compared to WT vacuoles, *npc2*Δ shows a loss of ergosterol and all three CSLs. (**C**) In the whole cell lipidome of early stationary stage cells, no reduction in ergosterol or SL abundance is observed for *npc2*Δ or *ncr1*Δ compared to WT. (**D**) Schematic for the yeast NPC system’s potential function in distributing lipid components from internalized cargoes. Classically, Npc2 shuttles sterols to the receptor Ncr1 for incorporation into the vacuole membrane (solid arrow). Trafficking of larger, polar lipids is not accommodated by Ncr1’s NTD, suggesting that Npc2 can act independently or through an alternative receptor (dashed arrow).

### Genetic modulation of CSL composition in the yeast lipidome

To directly test the hypothesis that changes in SL composition can drive vacuole phase separation, we generated a set of yeast strains to systematically modulate the abundance of each CSL in cells (Fig. 3A). The first phosphotransferase in the pathway, Aur1, transfers an inositol phosphate from PI to ceramide phytoceramide, generating IPC (Kuroda et al., 1999). *AUR1* is an essential gene, so we utilized a promoter replacement strategy to place its expression under control of the repressor doxycycline (P_tetoff_-*AUR1*). The subsequent glycosyltransferase and phosphotransferase to generate MIPC and M(IP)_2_C, respectively, are non-essential and so gene knockouts could be utilized to test the function of these species. Csg1, Csh1 and Csg2 are the subunits that form MIPC synthase (Uemura et al., 2003); the former two are paralogs that each form complexes with the latter. To create strains that lack MIPC synthesis, we generated strains lacking Csg1 and Csh1 (*csg1*Δ*csh1*Δ) or in Csg2 (*csg2*Δ). Synthesis of M(IP)_2_C occurs by a single gene product, Ipt1 (Dickson et al., 1997), so loss of M(IP)_2_C was assayed in *ipt1Δ*. The strains generated to analyze CSL effects on vacuole domain formation are summarized in Supplementary Table 1.

**Figure 3:**
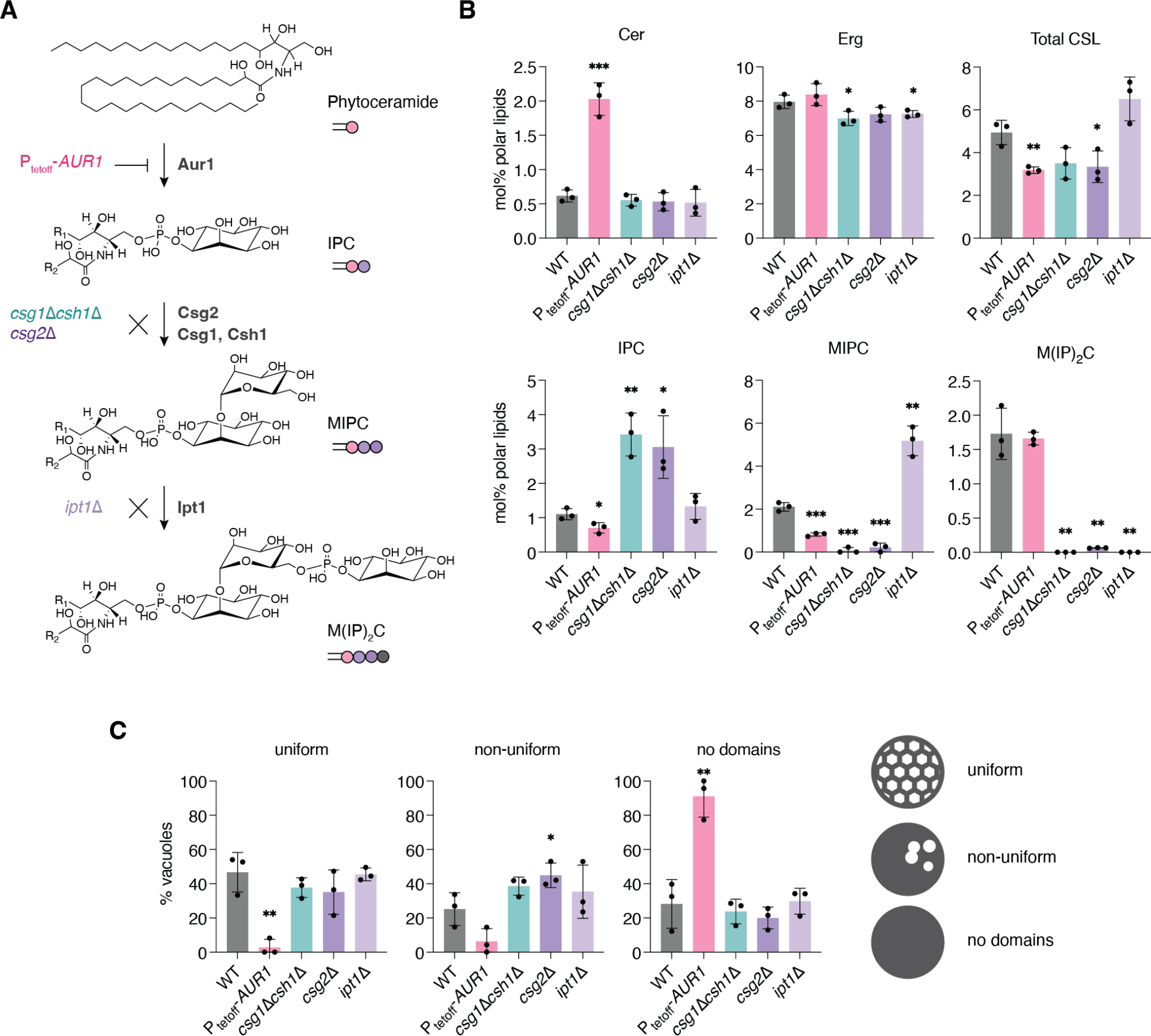
Genetic dissection of the CSL composition and effects on vacuole domains. (**A**) The biosynthetic pathway from phytoceramide to three abundant CSLs: IPC, MIPC, M(IP)_2_C. Successive action by the Aur1, the Csg2 with Csg1 or Csh1 complex, and Ipt1 produce the three abundant CSLs. To the left of each step are shown the strain modifications that manipulate them: knockdown of *AUR1* (P_tetoff_-*AUR1* grown in the presence of doxycycline), deletion of *CSG2* (*csg2*Δ) or *CSG1* and *CSH1* (*csg1*Δ*csh1*Δ), and deletion of *IPT1* (*ipt1*Δ). (**B**) Whole cell lipidomics of the strains indicated in panel A, showing the expected changes in the abundance of each SL, as well as total CSL and ergosterol abundances. Significance was assessed by unpaired two-tailed t-test against WT; *, p < 0.05; **, p < 0.01; ***, p < 0.001. (**C**) Resulting changes in vacuole domain frequency and type for each mutant. Examples of each domain type are shown in Fig. S5. Domain morphology classifications were quantified and plotted for the strains (n = 3 individual cultures, N > 100 cells for each). Significance was assessed by unpaired two-tailed t-test against the WT; *, p < 0.05; **, p < 0.01.

To verify that mutations altered the pathway as expected, we first analyzed the lipidomes of whole cells grown under domain-formation conditions (Fig. 3B). In general, lipid abundances changed as expected: P_tetoff_-*AUR1* cells grown in the presence of doxycycline (*AUR1* knockdown) showed a substantial (1.5-fold) decrease in total CSLs compared to wild-type (WT) cells and a corresponding increase in ceramides, the substrates for Aur1. Interestingly, the CSL pool in *AUR1* knockdown cells was dominated by M(IP)_2_C. Both *csg2*Δ and *csg1*Δ*csh1*Δ strains had very similar lipidomes containing low abundances amounts of MIPC and only trace amounts of M(IP)_2_C, with a corresponding increase in IPC compared to WT. As previously reported, *ipt1*Δ cells did not produce M(IP)_2_C and accumulated the precursor MIPC, with no changes to IPC levels. Subtle changes to phospholipid species were observed in response to CSL manipulation, suggesting compensatory changes in lipid metabolism (Fig. S4). Most notably, PI levels increased in P_tetoff_-*AUR1*, *csg2*Δ, or *csg1*Δ*csh1*Δ strains compared to WT at the expense of either PE (*AUR1* knockdown) or PC (*csg2*Δ, or *csg1*Δ*csh1*Δ). Because PI is a substrate for Aur1, its increase upon *AUR1* knockdown was expected, but the similarity of that effect to those in *csg2*Δ or *csg1*Δ*csh1*Δ, which do not accumulate IPC, suggest additional interactions between PI and yeast SL metabolism. Ergosterol levels showed only minor changes in these mutants.

### Vacuole CSL composition correlates with domain frequency and morphology

We next characterized the effects of CSL perturbation on the ability of cells to undergo vacuole phase separation (Fig. 3C). We classified cells as showing either 1) uniform Lo domains, forming a distinctive soccer ball-like pattern 2) non-uniform domains, which were generally smaller, irregularly spaced, and often aggregated or 3) no domains, characterized by an apparent homogeneous distribution of Pho8-GFP (Fig. S5). Quantification of large numbers of vacuoles from each strain (N>100 per replicate) showed that WT and *ipt1*Δ cells had similar vacuole morphologies, in which the majority of the vacuoles contained uniform domains. In contrast, *csg2*Δ and *csg1*Δ*csh1*Δ cells, which largely lack MIPC and M(IP)_2_C, exhibited vacuoles with non-uniform domains. Cells with reduced *AUR1* expression contained vacuoles that predominantly lacked domains. These results indicate that alteration of CSL metabolism is sufficient to modulate the morphology and frequency of vacuole domains.

Based on the results above, we isolated stationary stage vacuoles from a subset of our mutants to directly correlate changes in domain formation with the membrane lipidome. We compared early stationary stage vacuoles from WT cells with those from *AUR1* knockdown cells, featuring reduced domain frequency, and from *csg2*Δ cells, featuring altered domain morphology (Fig. 4A). *AUR1* knockdown vacuoles were characterized by a large (64.3%) reduction in total CSL content compared to WT vacuoles. Of the CSLs, IPC showed the largest reduction, but significant decreases were present in all species. In contrast, *csg2*Δ vacuoles showed an identical total CSL content as in WT, but an accumulation of IPC and almost complete loss of MIPC and M(IP)_2_C. Thus, both the abundances and distributions of CSLs in these vacuoles differed dramatically from each other (Fig. 4B). The vacuole lipidomes of SL mutants contained common signatures in other lipid classes when compared to WT vacuoles (Fig. S6): moderately lower ergosterol content (25% less than WT), an increase in the PC/PE ratio, and an increase in PS. The *AUR1* knockdown vacuoles also showed an increase in PI, a substrate for IPC synthase.

**Figure 4:**
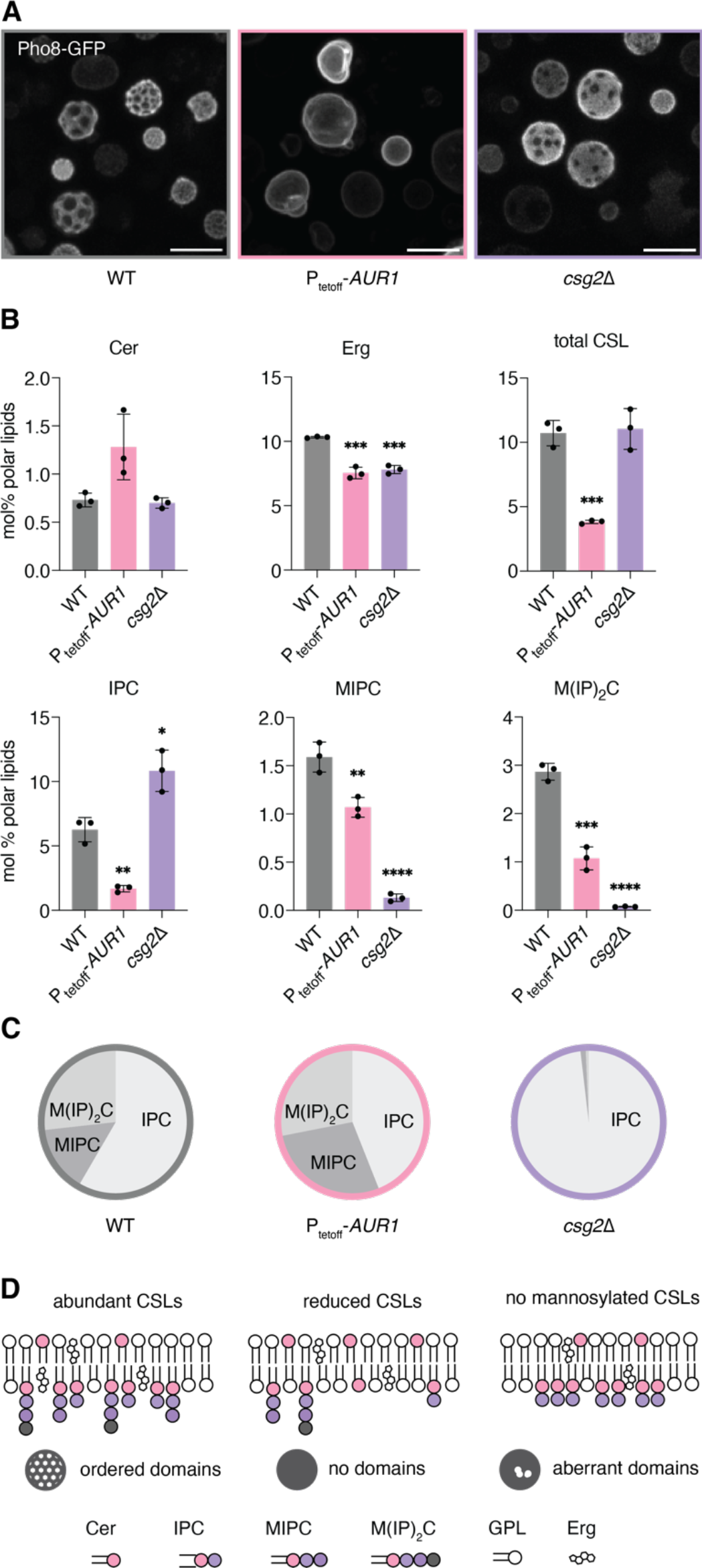
Correlating vacuole lipid composition with propensity for domain formation in CSL mutants. (**A**) A representative field view of WT, P_tetoff_-*AUR1* grown with doxycycline (*AUR1* knockdown) and *csg2*Δ cells, shown as 3D projections of confocal Z-stacks that were visualized using Ld vacuole domain marker, Pho8-GFP. Scale bars, 5 µm. (**B**) Vacuole lipidomics show changes to SLs and ergosterol levels in vacuoles purified from *AUR1* knockdown and *csg2*Δ cells. The total CSL abundance is significantly reduced in *AUR1* knockdown vacuoles, whereas no change was observed in *csg2*Δ compared to the WT. Significance was assessed by unpaired two-tailed t-test against the WT; *, p < 0.05; **, p < 0.01; ***, p < 0.001; ****, p < 0.0001. (**C**) In addition to the total amount, the distribution of SL headgroups changes in vacuoles isolated from these mutants, with the proportion of IPC mildly reduced and dramatically increased in *AUR1* knockdown and *csg2*Δ vacuoles, respectively. (**D**) Proposed model for how SL composition in the vacuole membrane alters its domain organization.

While the complexities of real cell membrane composition make direct comparisons of individual lipids challenging, the similar levels of ergosterol and PC/PE ratios of *AUR1* knockdown and *csg2*Δ vacuoles argue that CSL abundance (3-fold higher in the latter, Fig. 4C) directly contributes to the frequency of vacuole phase separation (4-fold higher in the latter). Similarly, the comparison between WT and *csg2*Δ vacuoles, which have identical levels of CSLs but different stoichiometries between them, suggests that head group chemistry controls the abundance of non-uniform membrane domains (2-fold higher frequency of cells with this phenotype in *csg2*Δ). Compared to the uniform domains observed in WT cells, the non-uniform *csg2*Δ domains generally occupied a lower surface area, were often irregularly shaped, and showed a low propensity to coarsen upon fusion with other domains they came in contact with (Fig. S7), indicating that they lack fluidity associated with Lo domains in WT vacuoles. These data suggest that abundant CSLs promote membrane domains but a balance between IPC and mannosylated species is required to prevent gel-like domains from forming (Fig. 4D).

### CSL metabolism controls micro-lipophagy under glucose restriction

Vacuole domains are required for long-term survival under glucose restriction (Seo et al., 2017; Tsuji et al., 2017), so we asked if there were differences between microautophagy-related phenotypes between *AUR1* knockdown, *csg2*Δ, and WT cells. We observed that LDs, labeled with Erg6-dsRed, dock to uniform vacuole domains, which are dominant in WT cells, but also to non-uniform domains, common in *csg2*Δ cells (Fig. 5A). In contrast, cells lacking vacuole domains, which are dominant in cells with reduced *AUR1* expression, featured LDs that remained peripheral to the vacuole, but never fully docked to its membrane. We assayed micro-lipophagy during starvation conditions by measuring the partial degradation of LD-associated Erg6-GFP into free GFP, which occurs by vacuole proteases upon LD internalization (Fig. 5B). In comparison to WT, *AUR1* knockdown cells showed low accumulation of free GFP. In contrast, *csg2*Δ cells showed no defects in LD degradation by this assay. These observations suggest that while vacuole domains, reduced in *AUR1* knockdown cells, are necessary for LD binding and internalization, their characteristic morphology, lost in *csg2*Δ cells, is not.

**Figure 5:**
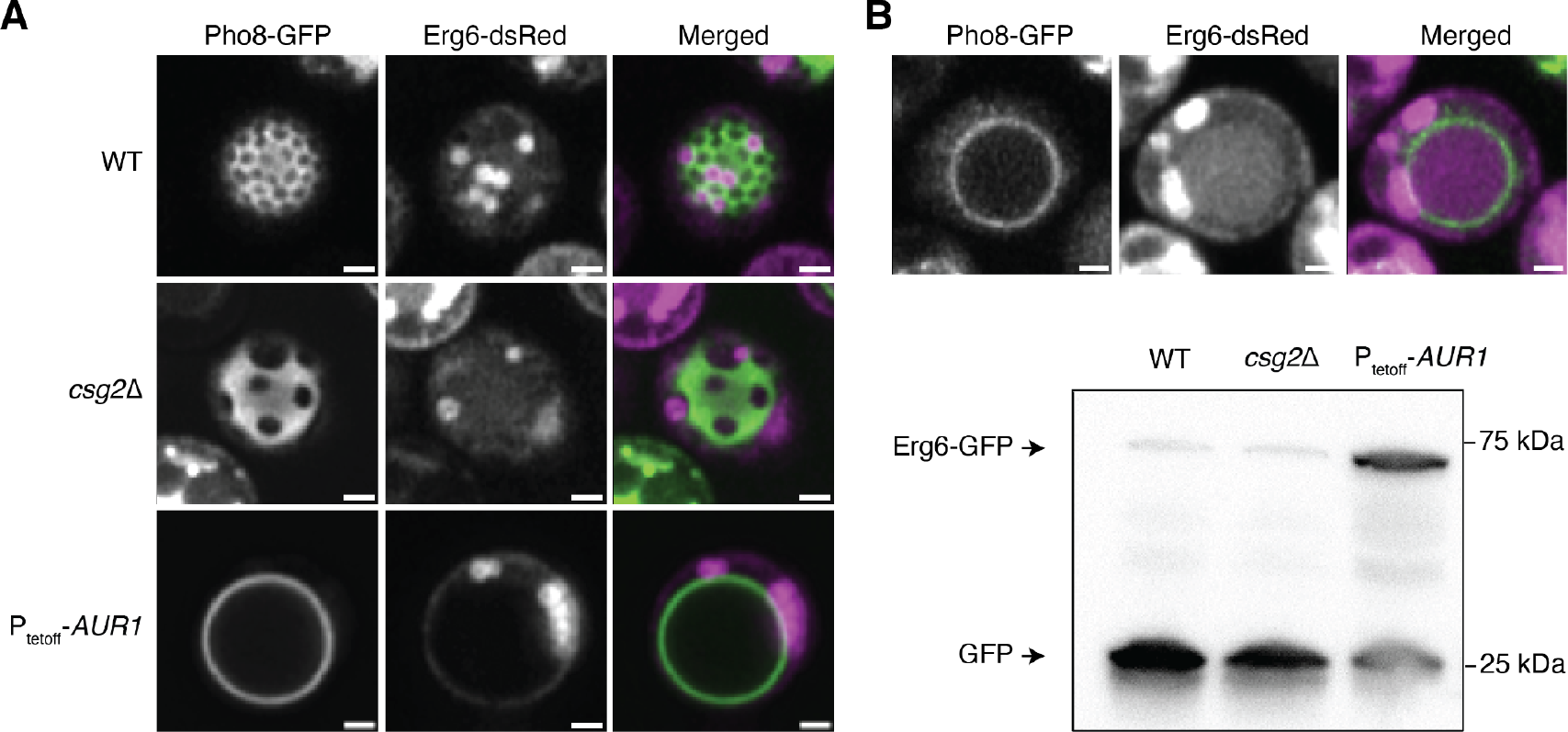
LD docking and micro-lipophagy is impaired by loss of CSLs. (**A**) LDs associate with both uniform and non-uniform Lo domains in WT and *csg2*Δ, cells, but fail to dock in P_tetoff_-*AUR1* cells grown with doxycycline (*AUR1* knockdown), which lack Lo domains. Scale bars, 1 µm. (**B**) Once LDs are docked and internalized, they are digested inside the vacuole. Shown above is a WT cell after 24 hours glucose restriction; the vacuole lumen contains degraded LDs, as indicated by the free dsRed signal inside of the vacuole. In the western blot below, intact LDs are measured in bulk by abundance full length Erg6-GFP, while free GFP is indicative of digestion of LDs and degradation of Erg6-GFP. Compared to WT and *csg2*Δ, more intact Erg6-GFP remains in *AUR1* knockdown cells.

## Discussion

An emerging model for vacuole organization is that autophagic lipid flux not only requires ordered membrane domains, which act as docking sites for LDs and other cargoes, but also promotes their formation through subsequent incorporation of raft-promoting lipids to the vacuole membrane (Wang et al., 2014; Seo et al., 2021). Here we provide the first detailed lipidomics data supporting this hypothesis, finding that both ergosterol and SLs re-distribute to the vacuole membrane at the onset of early stationary stage. Key to these data was the identification of growth conditions in which robust domain formation can be observed while the cell wall is still readily digestible for organelle purification. We compared vacuoles from these conditions to those isolated from late exponential stage, where vacuoles are of similar size and density but completely lack domains. Lipidomics analysis found that the switch to early stationary stage corresponds to an increase in raft-forming components, including ergosterol and all three yeast CSLs, a potential trigger for membrane phase separation. Notably, these components did not increase in the corresponding whole cell lipidome, indicating that they are re-directed into the vacuole.

Sorting of SLs, and to a lesser extent ergosterol, in the stationary stage vacuole membrane is dependent on Npc2, a soluble lipid transfer protein that resides in the vacuole lumen. While mammalian NPC2 is exclusively associated with sterol transport from the lysosomal lumen, recent biochemical analyses of yeast Npc2 suggest that it is more broadly involved in trafficking of polar lipids. Through specific polar interactions in its lipid binding site, Npc2 can bind several anionic GPLs, most notably PI, whose headgroup mimics that of IPC, suggesting it could have additional role in trafficking non-sterol lipids in the vacuole (Moesgaard et al., 2020). In contrast, the sterol binding pocket in the N-terminal domain (NTD) of Ncr1 is smaller than that of Npc2, much more closely resembling that of human NPC1, and is thus unlikely to bind large, polar lipids (Winkler et al., 2019). One possibility is that Npc2 could shuttle yeast polar lipids, including SLs, from lumenal cargoes to the vacuole membrane, either directly or through alternative receptors (Fig. 2D). Npc2 could also impact SL sorting through secondary consequences of its canonical function in ergosterol transport. We note that stationary stage vacuoles from *npc2*Δ still retain a higher abundance of raft-forming components than WT late exponential vacuole, suggesting the involvement of other sorting factors.

Our study is the second recent attempt to characterize the lipidome of stationary stage vacuoles. The other used a very different isolation technique (MemPrep) and did not show any increase in ergosterol content and only modest changes to the CSLs of stationary stage vacuoles. One source of variability is the difference in growth conditions between the studies. Here, phase separated vacuoles were isolated from early stationary stage cells, harvested after only 24 hours of growth on low glucose minimal media. In contrast, the MemPrep-isolated vacuoles were isolated from late stationary stage cells after 48 hours of growth in richer, complete media containing higher levels of glucose and amino acids. The yeast lipidome changes dramatically during the stationary stage between 24 and 48 hours, a difference that is likely even larger given the differences in growth medium. Notably, an increase in high melting temperature PC lipids observed in MemPrep-isolated vacuoles, but not in our early stationary stage vacuoles, have also been observed in whole cells sampled at 48 hours of growth, but not at 24 hours (Klose et al., 2012). While vacuole phase separation could be achieved through different mechanisms depending on the growth stage, its rapid onset over the course of just a few hours, which our study focuses on, likely necessitates intracellular sorting of lipids from existing pools.

We were surprised by the magnitude by which CSLs become enriched in early stationary stage vacuole membranes – 3-fold that of domain-free vacuoles harvested only hours previously. The composition of the CSL pool also changed in stationary stage vacuoles, shifting to one enriched in IPC with longer and more hydroxylated chains. These findings motivated us to focus on CSLs as modulators of membrane phase separation. We found that reduction of IPC and total CSL content in the vacuole, achieved through reduced expression of *AUR1*, strongly inhibited domain formation and its associated phenotypes of LD micro-lipophagy and survival in long-term starvation conditions. In contrast, an increase in vacuole IPC content, which is found in *csg2*Δ cells, did not change domain abundance, but instead altered their appearance and dynamics. The non-uniform morphologies of *csg2*Δ vacuole domains and their resistance to coarsening (Fig. S7) are consistent with rigid, gel-like membranes that can be promoted by SLs (Goñi and Alonso, 2009). Previous studies have suggested that gel-like domains are especially promoted by IPC: the high Laurdan General Polarization values of IPC/ergosterol mixtures (Klose et al., 2010) and direct observations of gel-like nanodomains on the yeast PM that are inhibited by mannosylation of IPC into MIPC and M(IP)_2_C (Aresta-Branco et al., 2011). Gel-like domains are also found to be predominant in ternary mixtures of GPLs, cholesterol, and glucosylceramide, the closest analogue to IPC in metazoans (Varela et al., 2016).

All three CSLs (IPC, MIPC, and M(IP)_2_C) are among the most abundant lipids in *S. cerevisiae* cells, but it is not yet known how their composition is regulated in different compartments and during different growth stages. The biophysical properties and dynamics of yeast CSLs have also not yet been extensively investigated, partially as a result of the lack of available synthetic versions for use in model membrane experiments. In experiments utilizing yeast-derived lipid mixtures, it has been observed that IPC specifically increases membrane ordering in liposomes, especially those containing ergosterol (Klose et al., 2010). It has also been shown that the ability of yeast lipid extracts to phase separate when reconstituted in GUVs is lost in *elo3*Δ cells that synthesize CSLs with shorter chains (Ejsing et al., 2009; Klose et al., 2010). Thus, multiple lines of data support the role of this lipid class in formation of ordered membrane domains. An open question is what biophysical constraints dictate the stoichiometry between the three yeast CSL classes, each of which features pronounced differences in head-group size, hydrogen-bonding capacity, and anionic charge.

Yeast CSLs bear structural similarities to glycosylated lipids in the other eukaryotic lineages, all of which have been implicated in inducing ordered membrane phases and potentially domains. Glycosyl inositol phosphorylceramides, the most abundant sphingolipids in plant cells, have been shown to promote membrane ordering in model membranes (Cassim et al., 2021; Yu and Klauda, 2021) and can cluster in nanodomains on the plant PM (Cacas et al., 2016). Mammalian glycosylated sphingolipids, gangliosides, are also potent nucleator membrane domains. Despite their different biochemistry and increased complexity, gangliosides share some common structural features with yeast CSLs, including variable combinations of both hexose subunits and charged groups (sialic acids or phosphates). Extensive experimental and modeling studies of gangliosides (Sarmento et al., 2021), and their hexosyl ceramide precursors (Varela et al., 2016), have highlighted the complex roles their head group composition can have on local membrane structure and formation of membrane domains *in vitro*. Understanding the ‘sugar-code’ by which diverse glycosylated lipids can influence membrane organization could be aided by similar approaches applied to fungal CSLs as yeast feature a tractable compartment for investigating structural effects on membrane organization.

## Acknowledgements

Arnold Seo provided technical assistance and reagents to initiate the project. Members of the Budin lab provided comments on the manuscript. The work was supported by the National Science Foundation (MCB-2046303).

## Author Contributions

H.K. carried out the research. I.B. supervised the research. All authors wrote the paper.

## Methods

### Yeast strains, plasmids, and media

*Saccharomyces cerevisiae* W303a was used as the base strain throughout the study. All strains generated are listed in Supplementary Table 1. Integrations were generated by PCR-based homologous recombination of PCR-amplified cassettes transformed via the lithium acetate method. The gene of interest was replaced by selection markers to generate knockout strains. For P_tetoff_-*AUR1* strain, a tetO_2_-CYC1 promoter and expression cassette for the tetracycline-controlled transactivator were amplified from plasmid PCM224 was substituted into the 500 bp upstream from the *AUR1* start codon. For vacuole imaging, yeast strains were transformed with plasmid pRS426 GFP-Pho8 that allows expression of Pho8-GFP. YPD, complete synthetic medium (CSM), and minimal medium were used to grow yeast cells. YPD medium (Fisher Scientific) contained yeast extract (10g/L), peptone (20g/L) and glucose (20g/L). CSM medium contained glucose (20g/L), ammonium sulfate (5g/L) and yeast nitrogen base (Dibco) (1.7g/L). For CSM medium, amino acids and nucleobases were supplemented using CSM powders (MP Biomedicals). In order to promote vacuole phase separation, the yeast were diluted into minimal medium that contains 0.4% glucose, ammonium sulfate, yeast nitrogen base and only essential amino/nucleic acids. Minimal medium contained glucose (Fisher Scientific) (4g/L), ammonium sulfate (Fisher Scientific) (5g/L), yeast nitrogen base without ammonium sulfate and amino acids (BD Difco) (1.7g/L), leucine (Alfa Aesar) (20 mg/L), histidine (Arcos Organics) (20 mg/L), tryptophan (Alfa Aesar) (20 mg/L), adenine (Alfa Aesar) (10 mg/L) and uracil (Alfa Aesar) (20 mg/L), as described by Sherman et al. (Sherman, 2002) Inhibition of *AUR1* expression was induced by addition of 10 mg/mL of doxycycline (Sigma-Aldrich) stock solution to a final concentration of 10 µg/mL when the culture was diluted into minimal medium. Doxycycline added to WT cells at this concentration showed no effects on vacuole morphology or phase separation.

### Vacuole purification by density centrifugation

Yeast strains expressing Pho8-GFP were used to confirm the growth stage and domain formation under confocal microscope before starting vacuole purification. An overnight culture in 5mL YPD was prepared and grown in a shaker for ∼18 hrs, the culture was diluted 1/100 into 42mL CSM - uracil in a 125mL flask. After ∼18 hr incubation, 13 OD units of yeast were diluted to 650mL minimal media in a 2L non-baffled flask shaking at 200 rpm. For yeast in the late exponential phase, the cultures were incubated for 17 hrs after dilution into minimal medium (OD ∼ 0.7). For early stationary phase vacuole samples, yeasts were harvested after 24 hrs of incubation (OD ∼ 1.2). The cells were harvested by benchtop centrifugation (3,000xg for 30 minutes), yielding 2-3g of wet weight (wwt) cell pellet.

Initial vacuole purification was done by a method expanded from Wiederhold et al. (Wiederhold et al., 2009) with several modifications. The cell pellet was first resuspended in 10mL of 100mM Tris-HCl, pH 9.5, then 100uL of 1M DTT (Fisher Bioreagents) was added. After incubation at 30 °C for 10 minutes, cells were centrifuged at 4,000xg for 5 min, washed with 10mL of water, washed with 10mL of 1.1M sorbitol (Thermo Scientific), then resuspended in 5mL of 1.1M sorbitol. To generate spheroplasts, 9mg Zymolyase 20T (Amsbio) per gram wwt cells was dissolved in 1mL of 1.1M sorbitol, then added to the suspension. When purifying vacuoles from exponential phase yeast, 5mg Zymolyase 20T per gram wwt cells was used instead. After one hour of incubation at 30 °C (with manual swirling every 15 minutes), all procedures were conducted on ice or at 4 °C. Spheroplasts were washed by centrifuging the spheroplasts through a layer of 7mL of 7.5% Ficoll 400 (VWR) in 1.1M sorbitol at 4,000xg for 20 min. The spheroplasts were resuspended in 10mL 12% Ficoll 400 in 10mM Tris-MES, pH 6.9. For lysis, the suspension was transferred to a dounce homogenizer (15mL, Wheaton) and homogenized by 15 strokes using a tight (“B”) pestle. 10.7mL suspension was transferred to a centrifuge tube (Beckman Coulter) with 5.3mL of 12% Ficoll 400 in 10mM Tris-MES was layered on top of it. Centrifugation was performed at 50,000xg for 35 min in a SW 32.1Ti rotor (Beckman Coulter). Crude vacuoles were isolated from the top layer and collected using a pipet (∼1 mL) and diluted in 5mL 12% Ficoll 400 in 10mM Tris-MES.

Crude vacuoles were further purified to generate pure samples, free of other contaminants, especially associated LDs. Crude vacuoles were first lightly homogenized in a dounce homogenizer by stroking 5 times with the B pestle. 5.3 mL of vacuole suspension was transferred to a centrifuge tube, then layered with 5.3mL 8% Ficoll 400 in 10mM Tris-MES and 5.3mL 4% Ficoll 400 in 10mM Tris-MES. The vacuoles were further separated from the mixture through density centrifugation at 50,000xg for 35 min in a SW 32.1Ti rotor. After centrifugation, the top layer was collected. In order to further purify the vacuole, we adapted a low-speed centrifugation protocol by Wiemken et al. (Wiemken et al., 1979). For this, vacuoles were diluted with 0.6M sorbitol in 5 mM PIPES-AMPD, pH 6.8. The vacuoles were then purified through sorbitol/sucrose gradient: 2 volumes (2-5 mL) of the diluted vacuoles were layered on top of 1 volume of 0.4 M sorbitol, 0.2 M sucrose (Thermo Scientific) in 5 mM PIPES-AMPD, pH 6.8, which was layered on top of 1 volume of 0.36 M sorbitol, 0.24 M sucrose in 5 mM PIPES-AMPD, pH 6.8 (Wiemken et al., 1979). After centrifugation at 3500xg for 30 min, vacuoles sedimented at the bottom of the tube. The vacuoles were resuspended in 0.6M sorbitol buffer. Vacuole integrity was assessed immediately by imaging under wide-field fluorescence and transmitted light microscopy (Thermo EVOS equipped with 63x Nikon oil immersion objective) after each purification. Successful preparations showed predominantly large particles (vacuoles) that were Pho8-GFP positive. These were then snap frozen and kept at -80 °C before analysis.

### Western blot analysis of vacuole purity and LD micro-lipophagy

The purity of vacuoles was assessed by means of western blot analysis. 20µg protein, as measured by BCA assay, was loaded for each sample. Primary antibodies against different organelle markers included anti-Cox4 (Abcam, ab110272), anti-Dpm1 (Abcam, ab113686), anti-Vph1 (Abcam, ab113683) and anti-Pho8 (Abcam, ab113688). The LD contamination in purified vacuole samples was tested by isolating vacuoles from WT expressing Erg6-dsRed as a LD marker, before doing western blot with anti-RFP (Rockland, 600-401-379) against Erg6-dsRed. For assaying LD degradation, 2.5 OD units of cells were collected, then proteins were extracted as described previously (Kushnirov, 2000). 10µL of the protein extract was loaded for each blot and anti-GFP (Invitrogen, GF28R) was used as the primary antibody. As a secondary antibody, either goat anti-mouse IgG (H+L), HRP conjugate (Invitrogen, 31430) or goat anti-Rabbit IgG (H&L), HRP conjugate (ImmunoReagents, Inc., GtxRb-003-EHRPX) was used.

### Lipid extraction and lipidomics of yeast cells and isolated vacuoles

Mass spectrometry-based lipid analysis was performed by Lipotype GmbH (Dresden, Germany) as described (Ejsing et al., 2009; Klose et al., 2012). Lipids were extracted using a two-step chloroform/methanol procedure (Ejsing et al., 2009). Samples were spiked with internal lipid standard mixture containing: CDP-DAG 17:0/18:1, CL 14:0/14:0/14:0/14:0, Cer 18:1;2/17:0, DAG 17:0/17:0, LPA 17:0, LPC 12:0, LPE 17:1, LPI 17:1, LPS 17:1, PA 17:0/14:1, PC 17:0/14:1, PE 17:0/14:1, PG 17:0/14:1, PI 17:0/14:1, PS 17:0/14:1, EE 13:0, TAG 17:0/17:0/17:0, stigmastatrienol, IPC 44:0;2 MIPC 44:0;2 MIPC and M(IP)_2_C 44:0;2. After extraction, the organic phase was transferred to an infusion plate and dried in a speed vacuum concentrator. 1^st^ step dry extract was re-suspended in 7.5 mM ammonium acetate in chloroform/methanol/propanol (1:2:4, V:V:V) and 2^nd^ step dry extract in 33% ethanol solution of methylamine in chloroform/methanol (0.003:5:1; V:V:V). All liquid handling steps were performed using Hamilton Robotics STARlet robotic platform with the Anti Droplet Control feature for organic solvents pipetting.

Lipid extracts were analyzed by direct infusion on a QExactive mass spectrometer (Thermo Scientific) equipped with a TriVersa NanoMate ion source (Advion Biosciences). Samples were analyzed in both positive and negative ion modes with a resolution of R_m/z=200_=280000 for MS and R_m/z=200_=17500 for MSMS experiments, in a single acquisition. MSMS was triggered by an inclusion list encompassing corresponding MS mass ranges scanned in 1 Da increments(Surma et al., 2015). Both MS and MSMS data were combined to monitor EE, DAG and TAG ions as ammonium adducts; PC as an acetate adduct; and CL, PA, PE, PG, PI and PS as deprotonated anions. MS only was used to monitor CDP-DAG, LPA, LPE, LPI, LPS, IPC, MIPC, M(IP)_2_C as deprotonated anions; Cer and LPC as acetate adducts and ergosterol as protonated ion of an acetylated derivative (Liebisch et al., 2006).

Lipidomics data were analyzed with in-house developed lipid identification software based on LipidXplorer (Herzog et al., 2011, 2012). Data post-processing and normalization were performed using an in-house developed data management system. Only lipid identifications with a signal-to-noise ratio >5, and a signal intensity 5-fold higher than in corresponding blank samples were considered for further data analysis. Abundances for each lipid species, expressed as mol %, were compiled and are available in a spreadsheet as part of the Supplementary Information. Subsequent plotting and statistical tests were performed in Graphpad Prism.

### Quantitative analysis of phase separation in SL mutants

Yeast strains expressing Pho8-GFP, a Ld domain marker (Seo et al., 2017), were used for fluorescence microscopy. For each biological replicate, an overnight culture was first grown in 5mL YPD, then diluted 1/100 to 5mL CSM medium lacking uracil for selection and incubated for ∼18 hours. 0.1 OD unit of the CSM culture was diluted in 5mL minimal media and incubated for 24 hours prior to imaging. All incubations were done in 30 °C round bottom tubes (Fisher Scientific) shaking at 250 rpm. Prior to imaging, live yeast cells were immobilized in 8-well microscope chamber slides (Nunc Lab-tek, Thermo Fisher Scientific) pre-coated with 1-2 mg/mL concanavalin-A (MP Biomedicals). Vacuole images were acquired on a Zeiss LSM 880 confocal microscope with an Airyscan detector. Samples were imaged at room temperature using a Plan-Apochromat 63x/1.4 Oil DIC M27 objective. Excitation was through a 488 nm Argon laser set at 2% power. For each sample, two to four ∼6-8µm Z-stack images were acquired depending on the cell density, and Airyscan processed in ZEN Black using default settings. For image analysis, cells were categorized into 3 groups based on the vacuole domain morphology. Only vacuoles captured from top to bottom were used for the quantification, and at least 100 cells were quantified in a given sample. 3D projections were generated using the Z project tool in ImageJ.

### Statistical analysis

All experiments were performed in biological replicates grown from individual yeast colonies. Values are expressed as mean ± standard deviation. Significance analysis was determined by unpaired, two-tailed t-tests using the GraphPad Prism 9.

## Data Availability

The data that support this study are available within the paper and in the Supplementary Information files. Source data are provided with this paper.

## Supplementary Information

**Figure S1:**
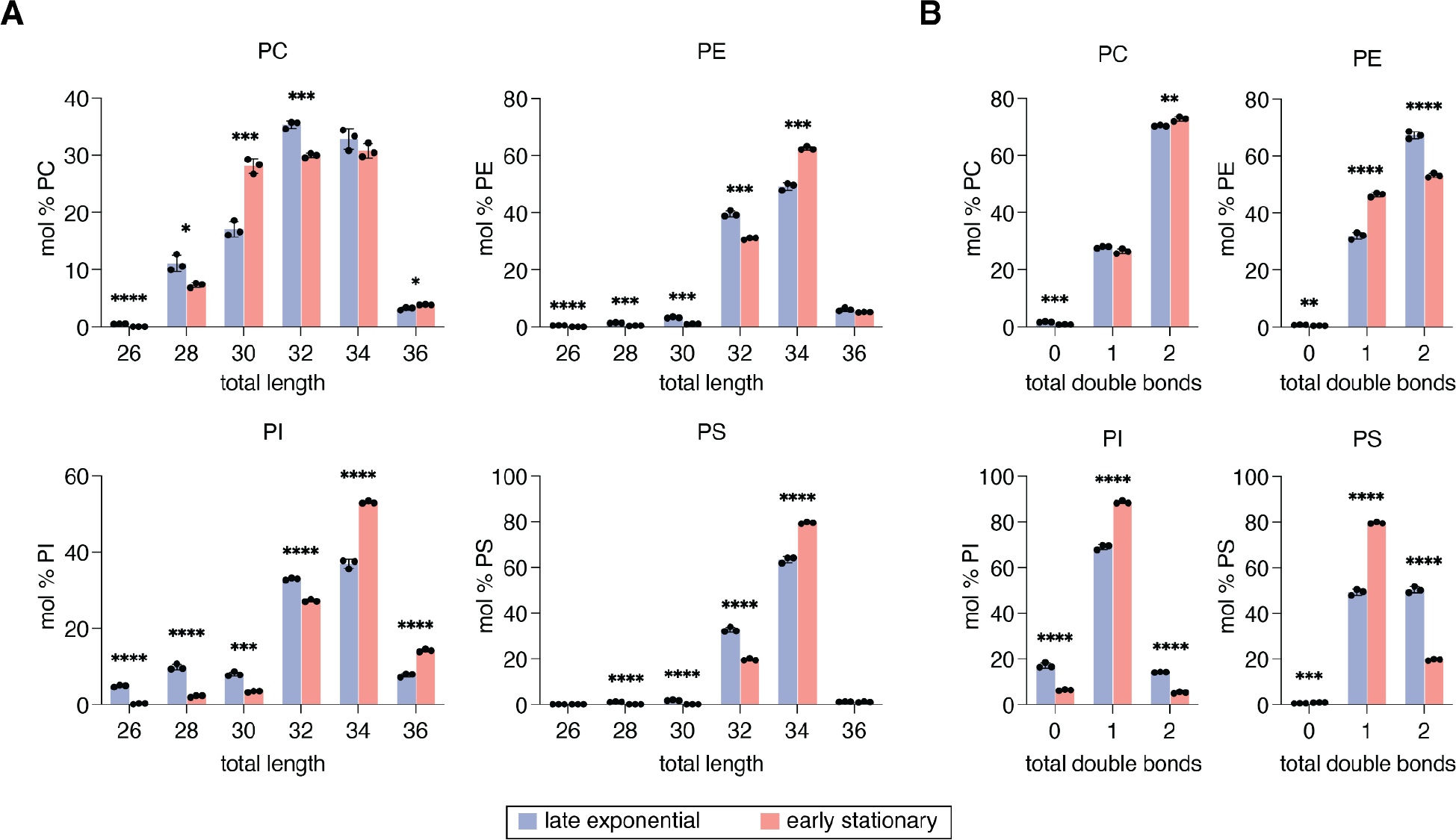
Molecular features of GPLs in isolated vacuoles. (**A**) Total acyl chain length, combining both acyl chains in PC, PE, PI, and PS of late exponential and early stationary stage vacuoles. PA and PG are not shown due to low abundance. (**B**) Number of double bonds (unsaturations) in late-exponential and stationary stage vacuoles. The proportion of monounsaturated PE, PI and PS lipids increases in the early-stationary stage, accompanied by a large decrease of di-unsaturated species. Significance was assessed by unpaired two-tailed t-test; *, p < 0.05; **, p < 0.01; ***, p < 0.001; ****, p < 0.0001.

**Figure S2:**
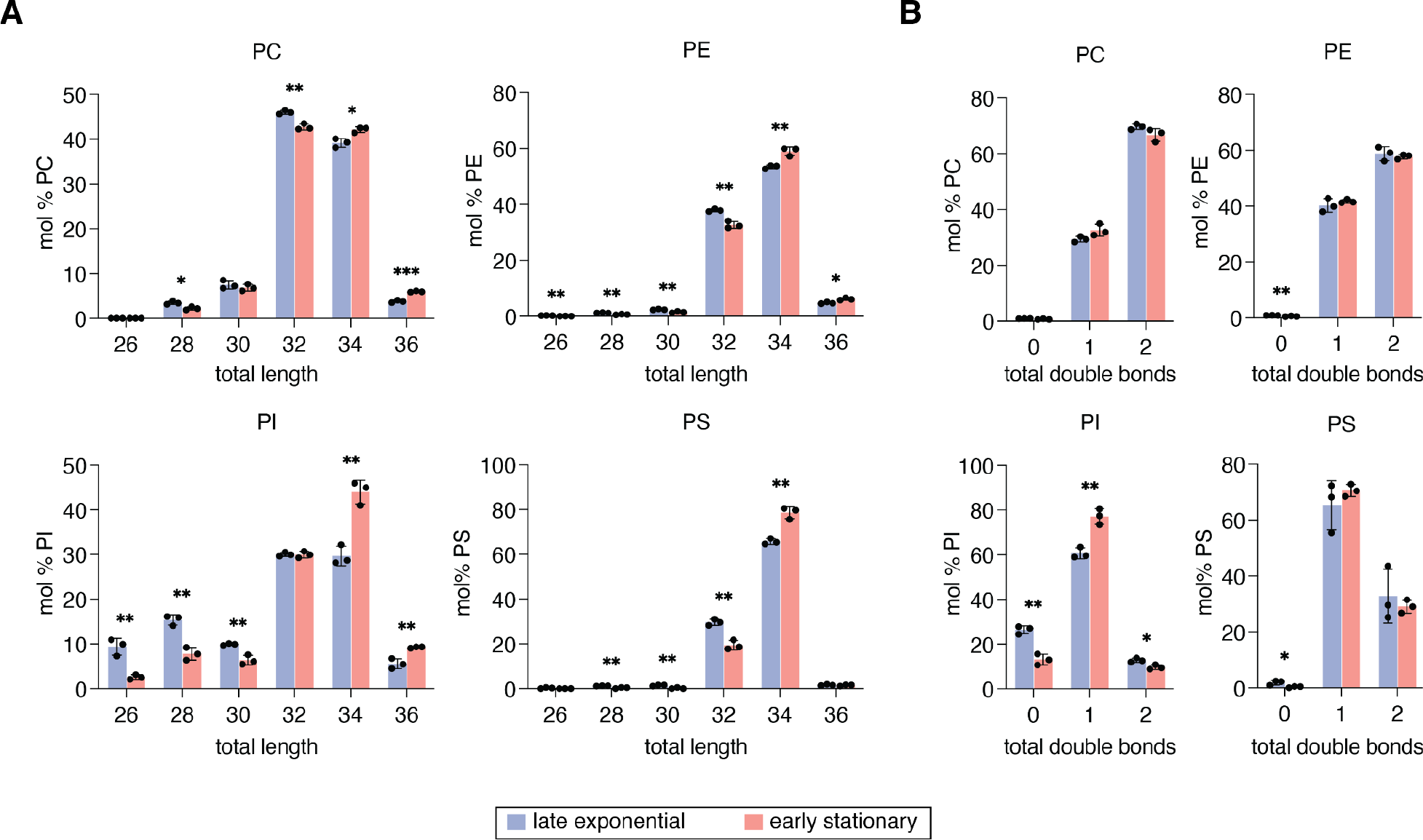
Molecular features of GPLs in whole cell lipid extracts. (**A**) Total acyl chain length, combining both acyl chains in PC, PE, PI, PS of the whole cell. The abundance of GPLs containing longer acyl chains (> 32 C) slightly increases in the stationary phase. (**B**) Total number of double bonds in PC, PE, PI and PS in whole cell lipidome. The unsaturation in PI is affected the most during the growth stage shift from late-exponential to early-stationary phase. The proportion of PI with only one double bond significantly increases, accompanied by decrease in PI with zero or two double bonds. Significance was assessed by unpaired two-tailed t-test; *, p < 0.05; **, p < 0.01; ***, p < 0.001.

**Figure S3:**
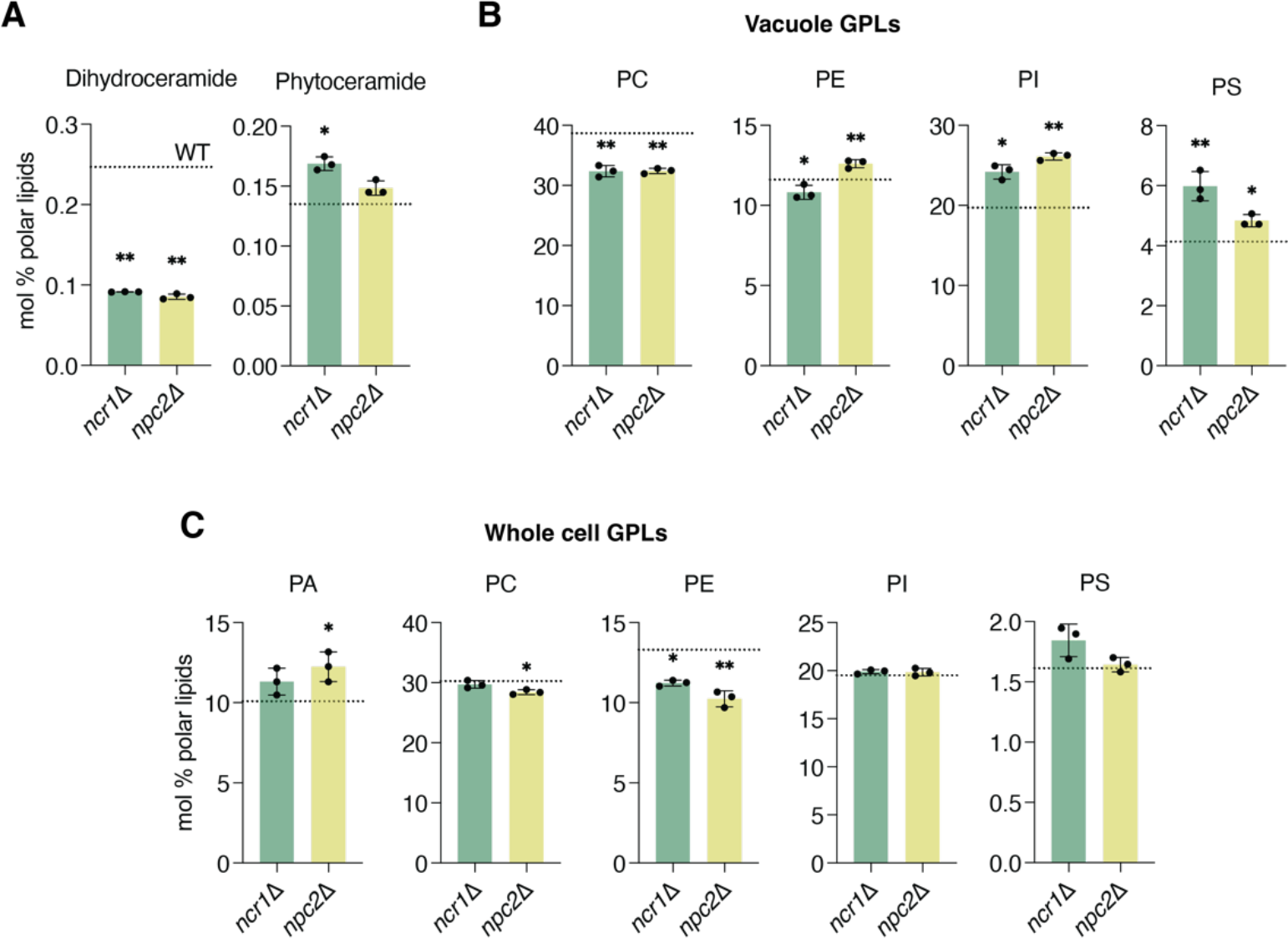
Changes in lipidome of *ncr1*Δ and *npc2*Δ early stationary stage cells and vacuoles, compared to WT cells (indicated by a dashed line in all panels). (**A**) The *ncr1*Δ whole cell lipidome shows decreases in dihydroceramides and increases in phytoceramides, consistent with previous analyses (Vilaça et al., 2018). (**B**) GPL levels in the vacuoles of *ncr1*Δ and *npc2*Δ show an increase in PI and loss of PC compared to WT. (**C**) GPL levels in the whole cells of *ncr1*Δ and *npc2*Δ show reduction in abundances of PE compared to WT. Significance was assessed by unpaired two-tailed t-test against the WT; *, p < 0.05; **, p < 0.01; ***, p < 0.001, ****, p < 0.0001.

**Figure S4:**
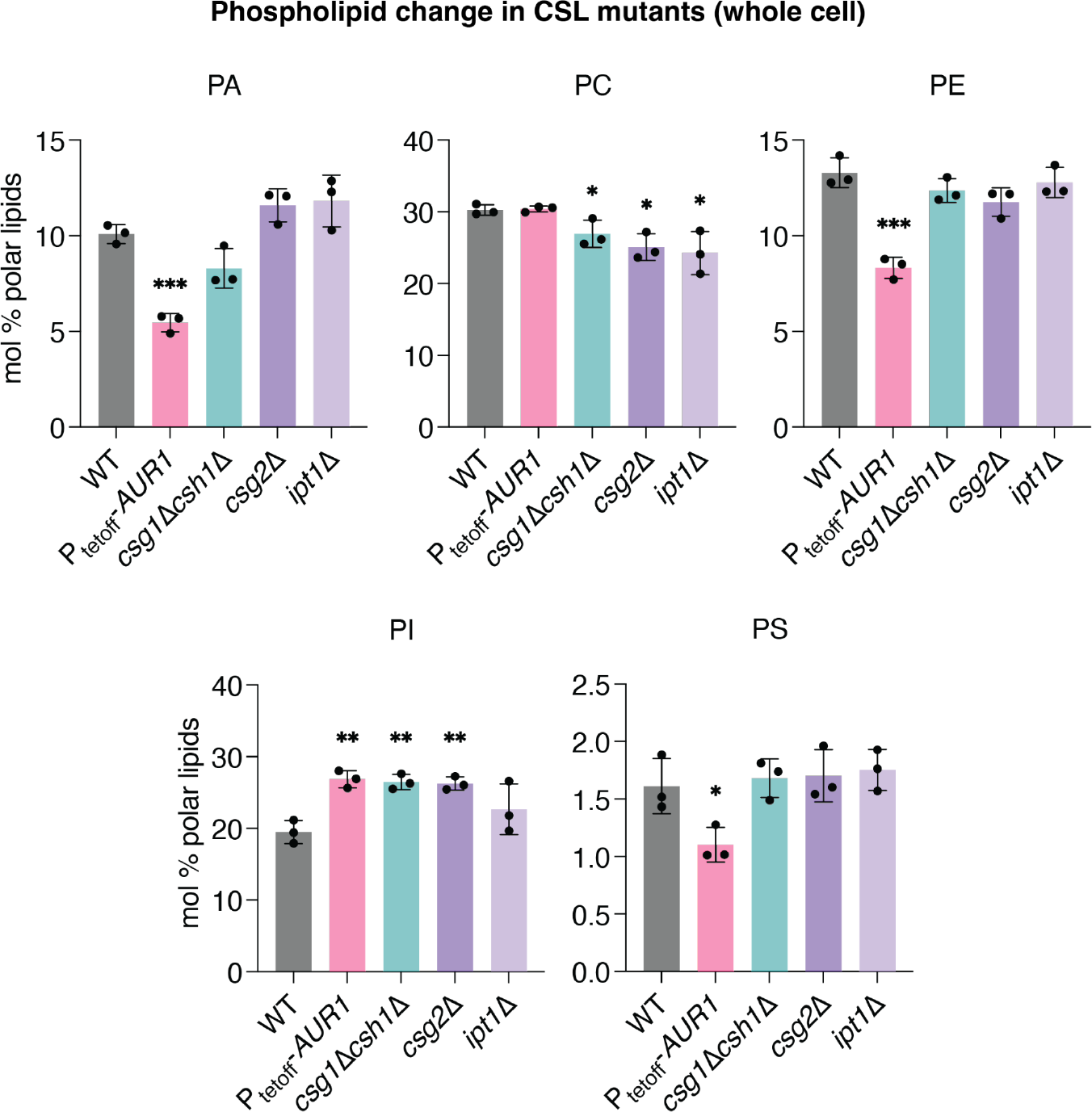
The change in different GPL classes in the whole cell lipidome of CSL mutants. The GPL profile of P_tetoff_-*AUR1* grown with doxycycline was significantly altered in all GPL classes except PC. The whole cell samples of *csg1Δcsh1Δ* and *csg2Δ* showed very similar changes, a decrease in PC and increase in PI compared to WT. Significance was assessed by unpaired two-tailed t-test against the WT; *, p < 0.05; **, p < 0.01; ***, p < 0.001.

**Figure S5:**
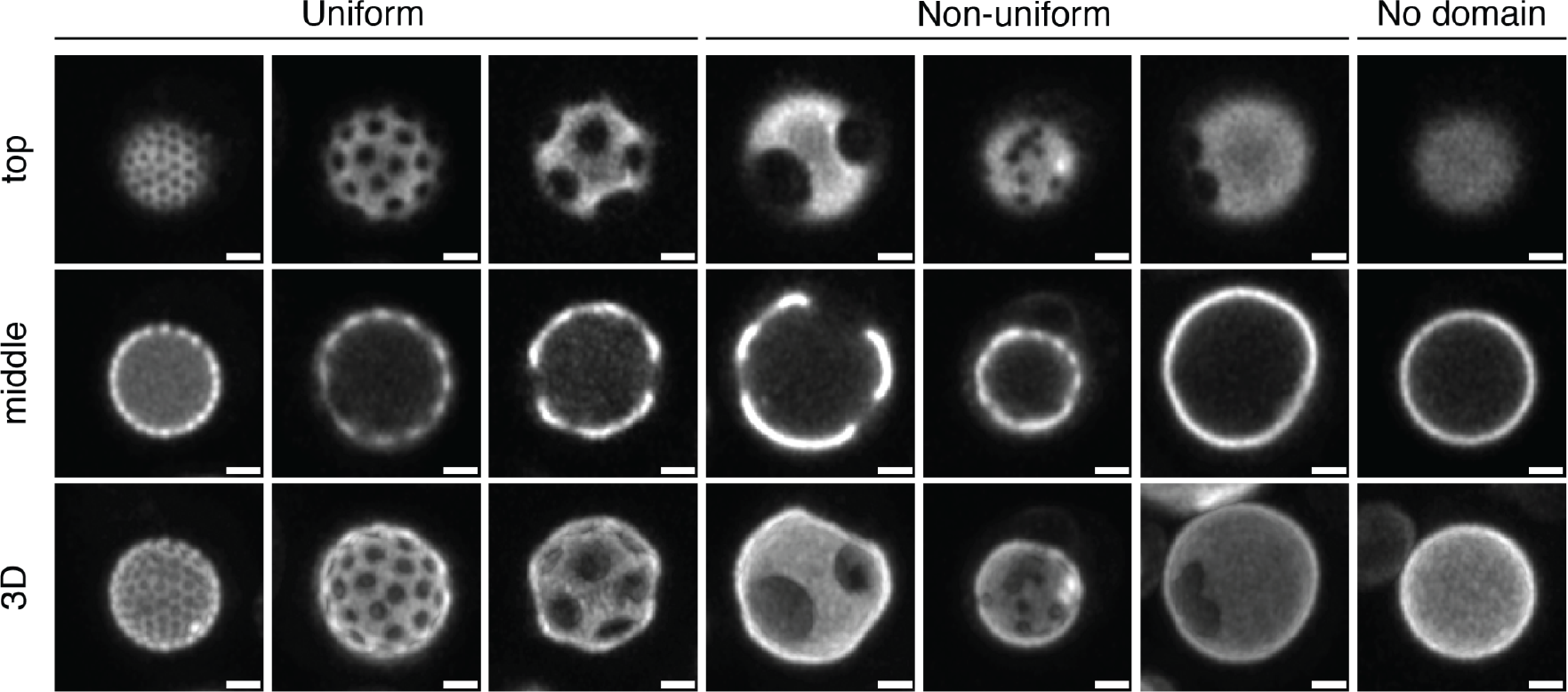
Categorization of vacuoles based on their domain morphology. Examples are shown of vacuole domain morphologies in early stationary stage cells. ‘Uniform’ vacuoles show domains that are uniformly distributed on the vacuole membrane, ‘non-uniform’ vacuoles have domains that are irregularly spaced out and ‘no domain’ vacuoles contain no observable vacuole microdomains under our imaging conditions. Shown are top and middle slices from typical Z-stacks and the resulting 3D projection. Scale bars, 1 µm.

**Figure S6:**
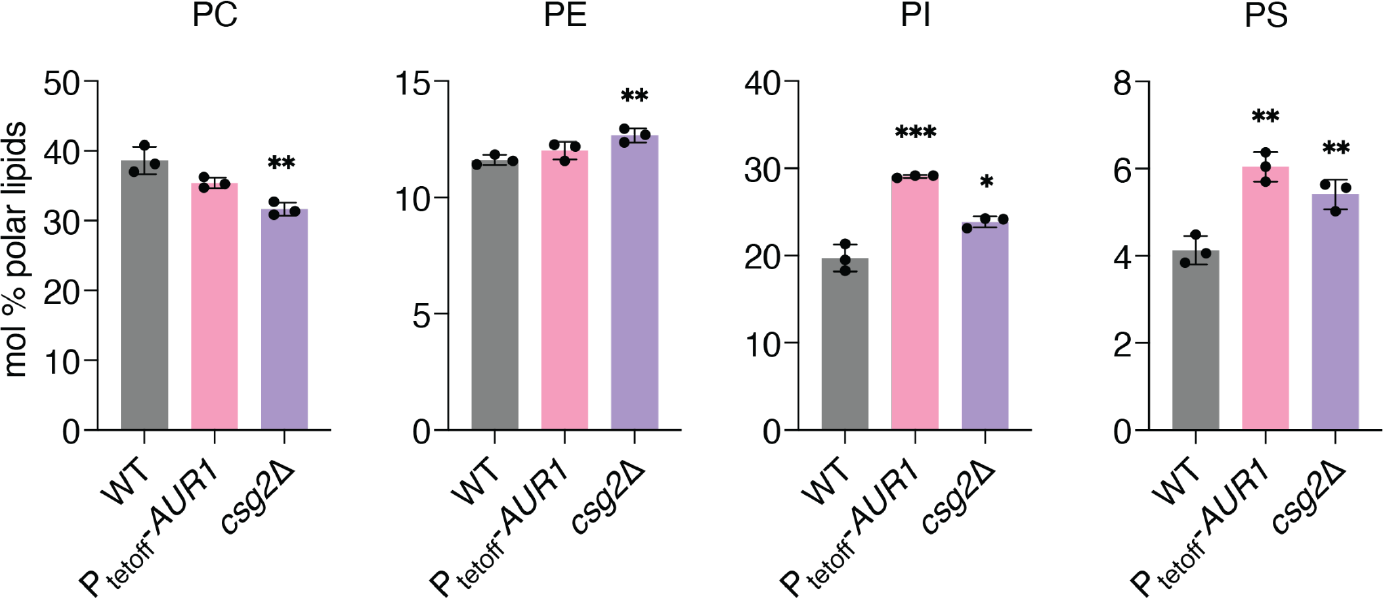
The change in different phospholipids in isolated vacuoles from P_tetoff_-*AUR1* cells (grown with doxycycline) and *csg2*Δ cells, compared to WT. The change does not follow the trend observed in the whole cell lipidome, except PC and PI. PE increases in *csg2*Δ vacuole, but not the whole cell. PE level is essentially the same in P_tetoff_-*AUR1*, when it was significantly reduced in the whole cell. The increase in PS was observed in the vacuole, which was not the case in the whole cell lipidome. Significance was assessed by unpaired two-tailed t-test against the WT; *, p < 0.05; **, p < 0.01; ***, p < 0.001.

**Figure S7:**
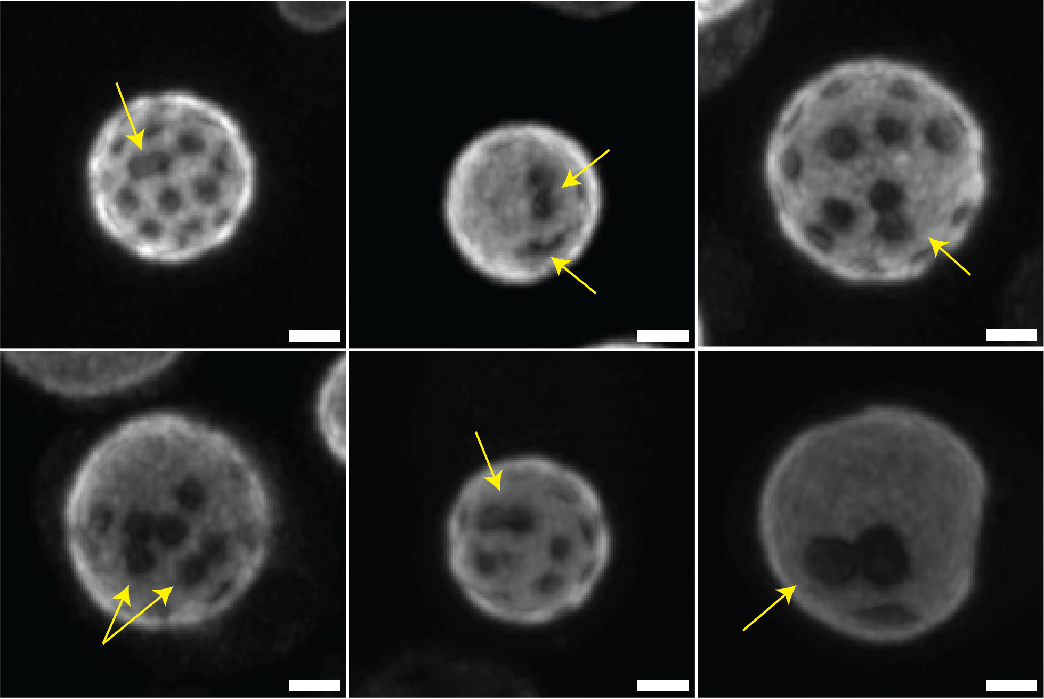
Vacuole domains do not readily coalesce upon collision in *csg2*Δ vacuoles. When liquid Lo domains typically coalesce, they form larger domains that quickly reform to minimize line tension(Rayermann et al., 2017). Shown are several examples where *csg2*Δ vacuoles showed ordered domains, imaged using Pho8-GFP (Ld marker), that upon fusion do not show rapid line tension minimization and retain their original shape (arrows). This could be explained by solid-like domains in *csg2*Δ vacuoles, in contrast to the liquid Lo domains typically observed in WT vacuoles. Scale bars, 1 µm.

**Table S1:** List of yeast strains used in this study.

| Strain | Genotype |
| --- | --- |
| W303a | *MATa ura3-52 trp1Δ2 leu2-3_112 his3-11*<br>*ade2-1 can1-100* |
| $P_{\text{tetoff}}\text{-}AUR1$ | W303a, *aur1::tetO<sub>2</sub>-P<sub>CYC1</sub>-AUR1* |
| *csg2Δ* | W303a, *csg2Δ::kanMX4* |
| *csg1Δcsh1Δ* | W303a, *csg1Δ::his6MX csh1Δ::LEU2* |
| *ipt1Δ* | W303a, *ipt1Δ::kanMX4* |
| *ncr1Δ* | W303a, *ncr1Δ::kanMX4* |
| *npc2Δ* | W303a, *npc2Δ::his6MX* |

## Notes

### Competing Interest Statement

The authors have declared no competing interest.

### Summary of Updates

Figures and text slightly updated.

## References

Aresta-Branco, F., A.M. Cordeiro, H.S. Marinho, L. Cyrne, F. Antunes, and R.F.M. de Almeida. 2011. Gel domains in the plasma membrane of Saccharomyces cerevisiae: highly ordered, ergosterol-free, and sphingolipid-enriched lipid rafts. J. Biol. Chem. 286:5043–5054.

Cacas, J.-L., C. Buré, K. Grosjean, P. Gerbeau-Pissot, J. Lherminier, Y. Rombouts, E. Maes, C. Bossard, J. Gronnier, F. Furt, L. Fouillen, V. Germain, E. Bayer, S. Cluzet, F. Robert, J.-M. Schmitter, M. Deleu, L. Lins, F. Simon-Plas, and S. Mongrand. 2016. Revisiting Plant Plasma Membrane Lipids in Tobacco: A Focus on Sphingolipids. Plant Physiol. 170:367– 384.

Cassim, A.M., Y. Navon, Y. Gao, M. Decossas, L. Fouillen, A. Grélard, M. Nagano, O. Lambert, D. Bahammou, P. Van Delft, L. Maneta-Peyret, F. Simon-Plas, L. Heux, B. Jean, G. Fragneto, J.C. Mortimer, M. Deleu, L. Lins, and S. Mongrand. 2021. Biophysical analysis of the plant-specific GIPC sphingolipids reveals multiple modes of membrane regulation. J. Biol. Chem. 296. doi:10.1016/j.jbc.2021.100602.

Dickson, R.C., E.E. Nagiec, G.B. Wells, M.M. Nagiec, and R.L. Lester. 1997. Synthesis of mannose-(inositol-P) 2-ceramide, the major sphingolipid in Saccharomyces cerevisiae, requires the IPT1 (YDR072c) gene. J. Biol. Chem. 272:29620–29625.

Ejsing, C.S., J.L. Sampaio, V. Surendranath, E. Duchoslav, K. Ekroos, R.W. Klemm, K. Simons, and A. Shevchenko. 2009. Global analysis of the yeast lipidome by quantitative shotgun mass spectrometry. Proc. Natl. Acad. Sci. U. S. A. 106:2136–2141.

Funato, K., and H. Riezman. 2001. Vesicular and nonvesicular transport of ceramide from ER to the Golgi apparatus in yeast. J. Cell Biol. 155:949–959.

Goñi, F.M., and A. Alonso. 2009. Effects of ceramide and other simple sphingolipids on membrane lateral structure. Biochim. Biophys. Acta. 1788:169–177.

Hancock, J.F. 2006. Lipid rafts: contentious only from simplistic standpoints. Nat. Rev. Mol. Cell Biol. 7:456–462.

Herzog, R., K. Schuhmann, D. Schwudke, J.L. Sampaio, S.R. Bornstein, M. Schroeder, and A. Shevchenko. 2012. LipidXplorer: a software for consensual cross-platform lipidomics. PLoS One. 7:e29851.

Herzog, R., D. Schwudke, K. Schuhmann, J.L. Sampaio, S.R. Bornstein, M. Schroeder, and A. Shevchenko. 2011. A novel informatics concept for high-throughput shotgun lipidomics based on the molecular fragmentation query language. Genome Biol. 12:R8.

Klose, C., C.S. Ejsing, A.J. García-Sáez, H.-J. Kaiser, J.L. Sampaio, M.A. Surma, A. Shevchenko, P. Schwille, and K. Simons. 2010. Yeast lipids can phase-separate into micrometer-scale membrane domains. J. Biol. Chem. 285:30224–30232.

Klose, C., M.A. Surma, M.J. Gerl, F. Meyenhofer, A. Shevchenko, and K. Simons. 2012. Flexibility of a Eukaryotic Lipidome – Insights from Yeast Lipidomics. PLoS ONE. 7:e35063. doi:10.1371/journal.pone.0035063.

Kuroda, M., T. Hashida-Okado, R. Yasumoto, K. Gomi, I. Kato, and K. Takesako. 1999. An aureobasidin A resistance gene isolated from Aspergillus is a homolog of yeast AUR1, a gene responsible for inositol phosphorylceramide (IPC) synthase activity. Mol. Gen. Genet. 261:290–296.

Kushnirov, V.V. 2000. Rapid and reliable protein extraction from yeast. Yeast. 16:857–860.

Leber, R., E. Zinser, G. Zellnig, F. Paltauf, and G. Daum. 1994. Characterization of lipid particles of the yeast, Saccharomyces cerevisiae. Yeast. 10:1421–1428.

Leveille, C.L., C.E. Cornell, A.J. Merz, and S.L. Keller. 2022. Yeast cells actively tune their membranes to phase separate at temperatures that scale with growth temperatures. Proc. Natl. Acad. Sci. U. S. A. 119. doi:10.1073/pnas.2116007119.

Levental, I., K.R. Levental, and F.A. Heberle. 2020. Lipid Rafts: Controversies Resolved, Mysteries Remain. Trends Cell Biol. 30:341–353.

Levine, T.P., C.A. Wiggins, and S. Munro. 2000. Inositol phosphorylceramide synthase is located in the Golgi apparatus of Saccharomyces cerevisiae. Mol. Biol. Cell. 11:2267–2281.

Liebisch, G., M. Binder, R. Schifferer, T. Langmann, B. Schulz, and G. Schmitz. 2006. High throughput quantification of cholesterol and cholesteryl ester by electrospray ionization tandem mass spectrometry (ESI-MS/MS). Biochim. Biophys. Acta. 1761:121–128.

Moeller, C.H., J.B. Mudd, and W.W. Thomson. 1981. Lipid phase separations and intramembranous particle movements in the yeast tonoplast. Biochim. Biophys. Acta. 643:376–386.

Moeller, C.H., and W.W. Thomson. 1979. An ultrastructural study of the yeast tonoplast during the shift from exponential to stationary phase. J. Ultrastruct. Res. 68:28–37.

Moesgaard, L., D. Petersen, M. Szomek, P. Reinholdt, M.B.L. Winkler, K.M. Frain, P. Müller, B.P. Pedersen, J. Kongsted, and D. Wüstner. 2020. Mechanistic Insight into Lipid Binding to Yeast Niemann Pick Type C2 Protein. Biochemistry. 59:4407–4420.

Moor, H., and K. Mühlethaler. 1963. FINE STRUCTURE IN FROZEN-ETCHED YEAST CELLS. J. Cell Biol. 17:609–628.

Munro, S. 2003. Lipid rafts: elusive or illusive? Cell. 115:377–388.

Olson, D.K., F. Fröhlich, R.V. Farese Jr, and T.C. Walther. 2016. Taming the sphinx: Mechanisms of cellular sphingolipid homeostasis. Biochim. Biophys. Acta. 1861:784–792.

Owen, D.M., A. Magenau, D. Williamson, and K. Gaus. 2012. The lipid raft hypothesis revisited--new insights on raft composition and function from super-resolution fluorescence microscopy. Bioessays. 34:739–747.

Rahman, M.A., R. Kumar, E. Sanchez, and T.Y. Nazarko. 2021. Lipid Droplets and Their Autophagic Turnover via the Raft-Like Vacuolar Microdomains. Int. J. Mol. Sci. 22. doi:10.3390/ijms22158144.

Rambold, A.S., S. Cohen, and J. Lippincott-Schwartz. 2015. Fatty acid trafficking in starved cells: regulation by lipid droplet lipolysis, autophagy, and mitochondrial fusion dynamics. Dev. Cell. 32:678–692.

Rayermann, S.P., G.E. Rayermann, C.E. Cornell, A.J. Merz, and S.L. Keller. 2017. Hallmarks of Reversible Separation of Living, Unperturbed Cell Membranes into Two Liquid Phases. Biophys. J. 113:2425–2432.

Reinhard, J., C.L. Leveille, C.E. Cornell, A.J. Merz, C. Klose, R. Ernst, and S.L. Keller. 2023. Remodeling of yeast vacuole membrane lipidomes from the log (one phase) to stationary stage (two phases). Biophys. J. doi:10.1016/j.bpj.2023.01.009.

Reinhard, J., L. Starke, C. Klose, P. Haberkant, H. Hammarén, F. Stein, O. Klein, C. Berhorst, H. Stumpf, J.P. Sáenz, J. Hub, M. Schuldiner, and R. Ernst. 2022. A new technology for isolating organellar membranes provides fingerprints of lipid bilayer stress. bioRxiv. 2022.09.15.508072. doi:10.1101/2022.09.15.508072.

Sangiorgio, V., M. Pitto, P. Palestini, and M. Masserini. 2004. GPI-anchored proteins and lipid rafts. Ital. J. Biochem. 53:98–111.

Sarmento, M.J., M.C. Owen, J.C. Ricardo, B. Chmelová, D. Davidović, I. Mikhalyov, N. Gretskaya, M. Hof, M. Amaro, R. Vácha, and R. Šachl. 2021. The impact of the glycan headgroup on the nanoscopic segregation of gangliosides. Biophys. J. 120:5530–5543.

Schuck, S., M. Honsho, K. Ekroos, A. Shevchenko, and K. Simons. 2003. Resistance of cell membranes to different detergents. Proceedings of the National Academy of Sciences. 100:5795–5800.

Schulz, T.A., and W.A. Prinz. 2007. Sterol transport in yeast and the oxysterol binding protein homologue (OSH) family. Biochim. Biophys. Acta. 1771:769–780.

Seo, A.Y., P.-W. Lau, D. Feliciano, P. Sengupta, M.A.L. Gros, B. Cinquin, C.A. Larabell, and J. Lippincott-Schwartz. 2017. AMPK and vacuole-associated Atg14p orchestrate μ-lipophagy for energy production and long-term survival under glucose starvation. Elife. 6. doi:10.7554/eLife.21690.

Seo, A.Y., F. Sarkleti, I. Budin, C. Chang, C. King, S.-D. Kohlwein, P. Sengupta, and J. Lippincott-Schwartz. 2021. Vacuole phase-partitioning boosts mitochondria activity and cell lifespan through an inter-organelle lipid pipeline. bioRxiv. 2021.04.11.439383. doi:10.1101/2021.04.11.439383.

Shelby, S.A., I. Castello-Serrano, K.C. Wisser, I. Levental, and S.L. Veatch. 2023. Membrane phase separation drives responsive assembly of receptor signaling domains. Nat. Chem. Biol. 19:750–758.

Sherman, F. 2002. Getting started with yeast. *In* Methods in Enzymology. C. Guthrie and G.R. Fink, editors. Academic Press. 3–41.

Simons, K., and E. Ikonen. 1997. Functional rafts in cell membranes. Nature. 387:569–572.

Surma, M.A., R. Herzog, A. Vasilj, C. Klose, N. Christinat, D. Morin-Rivron, K. Simons, M. Masoodi, and J.L. Sampaio. 2015. An automated shotgun lipidomics platform for high throughput, comprehensive, and quantitative analysis of blood plasma intact lipids. Eur. J. Lipid Sci. Technol. 117:1540–1549.

Thumm, M. 2000. Structure and function of the yeast vacuole and its role in autophagy. Microsc. Res. Tech. 51:563–572.

Toulmay, A., and W.A. Prinz. 2013. Direct imaging reveals stable, micrometer-scale lipid domains that segregate proteins in live cells. J. Cell Biol. 202:35–44.

Tsuji, T., M. Fujimoto, T. Tatematsu, J. Cheng, M. Orii, S. Takatori, and T. Fujimoto. 2017. Niemann-Pick type C proteins promote microautophagy by expanding raft-like membrane domains in the yeast vacuole. Elife. 6. doi:10.7554/eLife.25960.

Tuller, G., T. Nemec, C. Hrastnik, and G. Daum. 1999. Lipid composition of subcellular membranes of an FY1679-derived haploid yeast wild-type strain grown on different carbon sources. Yeast. 15:1555–1564.

Uemura, S., A. Kihara, J.-I. Inokuchi, and Y. Igarashi. 2003. Csg1p and Newly Identified Csh1p Function in Mannosylinositol Phosphorylceramide Synthesis by Interacting with Csg2p*. J. Biol. Chem. 278:45049–45055.

Varela, A.R.P., A.S. Couto, A. Fedorov, A.H. Futerman, M. Prieto, and L.C. Silva. 2016. Glucosylceramide Reorganizes Cholesterol-Containing Domains in a Fluid Phospholipid Membrane. Biophys. J. 110:612–622.

Veatch, S.L., and S.L. Keller. 2003. Separation of liquid phases in giant vesicles of ternary mixtures of phospholipids and cholesterol. Biophys. J. 85:3074–3083.

Vilaça, R., I. Barros, N. Matmati, E. Silva, T. Martins, V. Teixeira, Y.A. Hannun, and V. Costa. 2018. The ceramide activated protein phosphatase Sit4 impairs sphingolipid dynamics, mitochondrial function and lifespan in a yeast model of Niemann-Pick type C1. Biochim. Biophys. Acta Mol. Basis Dis. 1864:79–88.

Wang, C.-W., Y.-H. Miao, and Y.-S. Chang. 2014. A sterol-enriched vacuolar microdomain mediates stationary phase lipophagy in budding yeast. J. Cell Biol. 206:357–366.

Wiederhold, E., T. Gandhi, H.P. Permentier, R. Breitling, B. Poolman, and D.J. Slotboom. 2009. The yeast vacuolar membrane proteome. Mol. Cell. Proteomics. 8:380–392.

Wiemken, A., M. Schellenberg, and K. Urech. 1979. Vacuoles: The sole compartments of digestive enzymes in yeast (Saccharomyces cerevisiae)? Arch. Microbiol. 123:23–35.

Winkler, M.B.L., R.T. Kidmose, M. Szomek, K. Thaysen, S. Rawson, S.P. Muench, D. Wüstner, and B.P. Pedersen. 2019. Structural Insight into Eukaryotic Sterol Transport through Niemann-Pick Type C Proteins. Cell. 179:485–497.e18.

Yu, Y., and J.B. Klauda. 2021. Symmetric and Asymmetric Models for the Arabidopsis thaliana Plasma Membrane: A Simulation Study. J. Phys. Chem. B. 125:11418–11431.

Zinser, E., and G. Daum. 1995. Isolation and biochemical characterization of organelles from the yeast, Saccharomyces cerevisiae. Yeast. 11:493–536.

